# The Dynamics of Cytoplasmic mRNA Metabolism

**DOI:** 10.1101/763599

**Authors:** Timothy J. Eisen, Stephen W. Eichhorn, Alexander O. Subtelny, Kathy S. Lin, Sean E. McGeary, Sumeet Gupta, David P. Bartel

## Abstract

For all but a few mRNAs, the dynamics of metabolism are unknown. Here, we developed an experimental and analytical framework for examining these dynamics for mRNAs from thousands of genes. mRNAs of mouse fibroblasts exit the nucleus with diverse intragenic and intergenic poly(A)-tail lengths. Once in the cytoplasm, they have a broad (1000-fold) range of deadenylation rate constants, which correspond to cytoplasmic lifetimes. Indeed, with few exceptions, degradation appears to occur primarily through deadenylation-linked mechanisms, with little contribution from either endonucleolytic cleavage or deadenylation-independent decapping. Most mRNA molecules degrade only after their tail lengths fall below 25 nt. Decay rate constants of short-tailed mRNAs vary broadly (1000-fold) and are more rapid for short-tailed mRNAs that had previously undergone more rapid deadenylation. This coupling helps clear rapidly deadenylated mRNAs, enabling the large range in deadenylation rate constants to impart a similarly large range in stabilities.

**Highlights:**

- mRNAs enter the cytoplasm with diverse intra- and intergenic lengths
- mRNA deadenylation rates span a 1000-fold range and correspond to mRNA half-lives
- After their tails become short, mRNAs decay at rates that span a 1000-fold range
- More rapidly deadenylated mRNAs decay more rapidly upon reaching short tail lengths

## Introduction

mRNAs corresponding to different genes are degraded at substantially different rates, with some mRNAs turning over in minutes and others persisting for days (Dölken et al., 2008). Different conditions or developmental contexts can modify these rates, resulting in the destabilization of previously stable mRNAs, or vice versa (Rabani et al., 2011). These rate changes influence the dynamics of mRNA accumulation and, ultimately, the steady-state abundance of mRNAs.

Many proteins that promote mammalian mRNA degradation also can recruit deadenylase complexes. These include Pumilio (Van Etten et al., 2012), SMG5/7 (Muhlemann and Lykke-Andersen, 2010), GW182 (Fabian et al., 2011), BTG/TOB factors (Mauxion et al., 2009), Roquin (Leppek et al., 2013), YTHDF2 (Du et al., 2016), and HuR, TTP, and other proteins that bind AU- and GU-rich elements (Vlasova-St Louis and Bohjanen, 2011; Fabian et al., 2013). That these diverse modifiers of mRNA stability converge on deadenylation suggests that differences in deadenylation rates might explain a substantial fraction of the variation observed in mRNA stability.

In the past, the dynamics of mRNA deadenylation have been examined on a gene-by-gene basis, involving pulsed expression and subsequent mRNA analysis using RNase H to cleave the mRNA and RNA blots to probe for the poly(A)-tailed 3′ fragment. Because this procedure has been performed for only a handful of cellular mRNAs in yeast (Decker and Parker, 1993; Muhlrad et al., 1994; Hilgers et al., 2006) and mammals (Mercer and Wake, 1985; Wilson and Treisman, 1988; Shyu et al., 1991; Chen and Shyu, 1995; Gowrishankar et al., 2005), some fundamental questions, including the extent to which a global relationship exists between deadenylation rate and mRNA stability, have remained unanswered.

Here, we developed experimental and analytical tools for the global analysis of tail-length dynamics. Applying these tools to the mRNAs of cultured mouse fibroblasts generated a unique resource of initial cytoplasmic tail lengths, deadenylation rates, and decay parameters for mRNAs of thousands of individual genes, which in turn provided fundamental insights into cytoplasmic mRNA metabolism.

## Results

### Global Profiling of Tail-Length Dynamics

Two high-throughput sequencing methods, each with distinct advantages, were initially developed to profile poly(A)-tail lengths. One of these is PAL-seq (poly(A)-tail-length profiling by sequencing), which reports the cleavage-and-polyadenylation site for each polyadenylated molecule (Subtelny et al., 2014), whereas the other is TAIL-seq, which can measure poly(A)-tails that have been terminally modified with non-adenosine residues (Chang et al., 2014; Lim et al., 2016). Here, we developed PAL-seq v2, which combines these advantages and has the further benefit over both previous methods of more robust compatibility with contemporary Illumina sequencing platforms (Figure S1).

To observe the tail-length dynamics of endogenous mRNAs, we employed a metabolic-labeling approach in which mRNAs of different age ranges were isolated and analyzed (Figure 1A). To initiate labeling, we added 5-ethynyl uridine (5EU) to 3T3 cells. After incubating for time periods ranging from 40 min to 8 h, cytoplasmically enriched lysates were collected, and RNA containing 5EU was isolated by virtue of the reactivity between the 5EU and a biotin-containing tag. Poly(A)-tail lengths of captured mRNAs, as well as total-lysate mRNA, were measured using PAL-seq v2 (hereafter called PAL-seq). In parallel, we performed RNA-seq, which provided measurements of abundance for mRNAs of each time interval. Spike-ins of RNA standards enabled estimates of recovery and measurement accuracy over a broad range of tail lengths, as well as absolute quantification of RNA measured by each method. These experiments were each performed using each of two independently passaged 3T3 cell lines. Unless stated otherwise, all figures show the results obtained for cell line 1. Nonetheless, the results of the two cell lines were highly reproducible at each time interval (*R*_s_ ≥ 0.81 for mean tail-length measurements). Moreover, results from either PAL-seq v1, PAL-seq v2, or our implementation of TAIL-seq were highly correlated (Figure S2A–D; *R*_s_ = 0.83–0.88 for each of the two-way comparisons), which indicated that our conclusions were independent of the method used for tail-length profiling.

**Figure 1.**
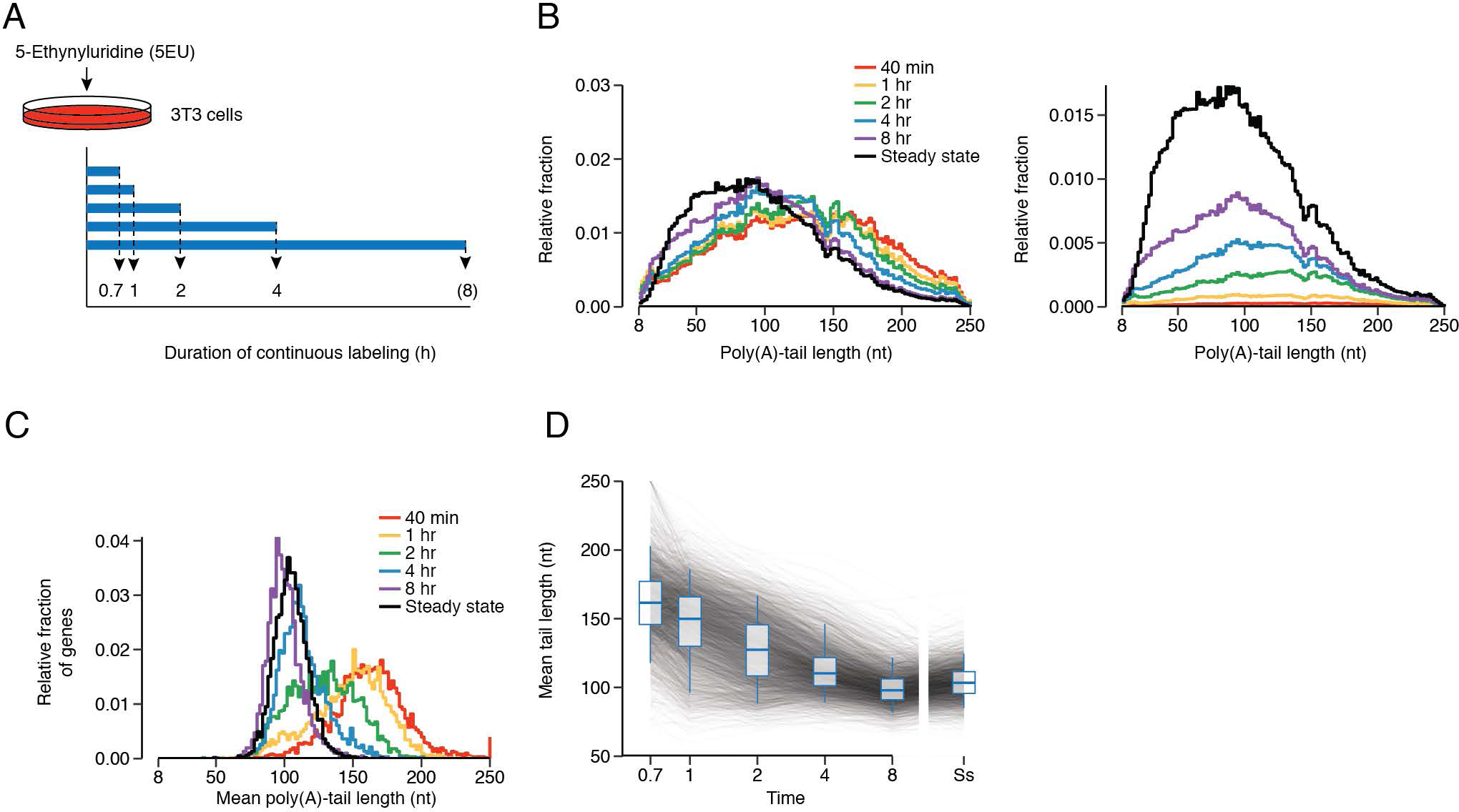
Global Tail-Length Dynamics of Mammalian mRNAs. (A) Schematic of 5EU metabolic-labeling. Experiments were performed with two 3T3 cell lines designed to induce either miR-155 or miR-1 (cell lines 1 and 2, respectively) but cultured without microRNA induction. The 8 is in parenthesis because an 8 h labeling period was included for only one line (cell line 1). For simplicity, all subsequent figures show the results for cell line 1, unless stated otherwise. (B) Tail-length distributions of mRNA molecules isolated after each period of 5EU labeling (key). Left: Distributions were normalized to each have the same area. Right: Distributions were scaled to the abundance of labeled RNA in each sample and then normalized such that the steady-state sample had an area of 1. The steady-state sample was prepared with unselected RNA from the 40 min time interval. Each bin is 2 nt; results for the bin with tail lengths ≥ 250 nt are not shown. (C) Distibutions of mean poly(A)-tail lengths for mRNAs of each gene after the indicated duration of 5EU labeling. Values for all genes that passed the tag cutoffs for tail-length measurement at all time intervals were included (n = 3048). Each bin is 2 nt. Genes with mean mRNA tail-length values greater than ≥ 250 nt were assigned to the 250 nt bin. (D) Tail lengths over time. Mean tail lengths for mRNAs from each gene (n = 3048) are plotted along with box-and-whiskers overlays (line, median; box, 25^th^ to 75^th^ percentiles; whiskers, 5^th^ to 95^th^ percentiles). See also Figures S1 and S2.

As expected if tail lengths become shorter over time in the cytoplasm (Sheiness and Darnell, 1973), mRNAs collected after the shortest labeling period (40 min) had the longest poly(A)-tail lengths, with a median length of 133 nt (Figure 1B). As the average age of each labeled mRNA population increased with longer labeling periods, tail-length distributions shifted towards the steady-state distribution (median length of 91 nt), with results from the 8 h period most closely resembling those of the steady state with respect to both length and abundance (Figure 1B). At each time interval, 10–20 nt tails preferentially possessed a 3′ terminal U (Figure S2E), although < 6.8% of tails had 3′ U residues in any sample, in keeping with previous reports on the fraction of short tails with terminal uridines at steady state (Chang et al., 2014; Lim et al., 2014). Analyses of mean poly(A)-tail lengths for mRNAs corresponding to thousands of individual genes showed that tails from mRNAs of essentially every gene shortened over time in the cytoplasm (Figure 1C–D).

### Correspondence Between mRNA Half-life and Deadenylation Rate

After 2 h of labeling, a broad range of mean tail lengths was observed, as mean tail lengths for mRNAs of some genes approached their steady-state values, whereas those for others still resembled their initial values (Figure 1C). These different rates of approach to steady-state tail lengths presumably at least partly reflected differences in mRNA degradation rates, as short-lived mRNAs were expected to reach their steady-state abundance and poly(A)-tail length more rapidly than were long-lived mRNAs.

To determine these degradation rates, we fit the yield of PAL-seq tags obtained for each gene at each time interval (normalizing to the spike-in controls) to the exponential function describing the approach to steady state, while also fitting a global offset to account for a delay between the time that 5EU was added and the time that labeled mRNAs appeared in the cytoplasm. This offset ranged from 27–36 min, depending on the experiment, a range consistent with single-gene measurements of the time required for mRNA transcription, processing, and export (Shav-Tal et al., 2004; Mor et al., 2010). Our half-life values (Table S1) correlated well with those previously reported for mRNAs of 3T3 cells growing in similar conditions (Schwanhäusser et al., 2011) (Figure S3A; *R*_s_ = 0.68–0.77), although our absolute values were substantially shorter (Figure S3B–D, median 2.1 h for mRNAs of the 3T3 cell line 2, as opposed to 9 h for previously reported values). This difference was attributable to potential divergence in the cell lines used in the two labs, as well as our focus on cytoplasmically enriched RNA and our absolute quantification of labeled RNA (made possible by adding standards to each sample prior to library preparation).

Previous analyses of the relationship between mRNA half-life and mean tail length have been limited to steady-state tail length measurements, for which no positive relationship has been observed (Subtelny et al., 2014), despite the established role of poly(A) tails in conferring mRNA stability. Our current datasets, which provided the opportunity make this comparison using half-life and tail-length measurements determined in the same study from the same cells, reinforced this finding; we observed no positive relationship between mRNA half-life and mean steady-state tail length (Figure S3G, *R*_s_ = –0.24). This result held when incorporating results of PAL-seq implemented with direct ligation to mRNA 3′ termini, which better detected very short or highly modified tails (Figure 2A, *R*_s_ = –0.02). Indeed, the mean tail lengths of long-lived mRNAs, including those of ribosome-protein genes (RPGs), closely resembled tail lengths of short-lived mRNAs, including those of immediate-early genes (IEGs) (Figure 2A).

**Figure 2.**
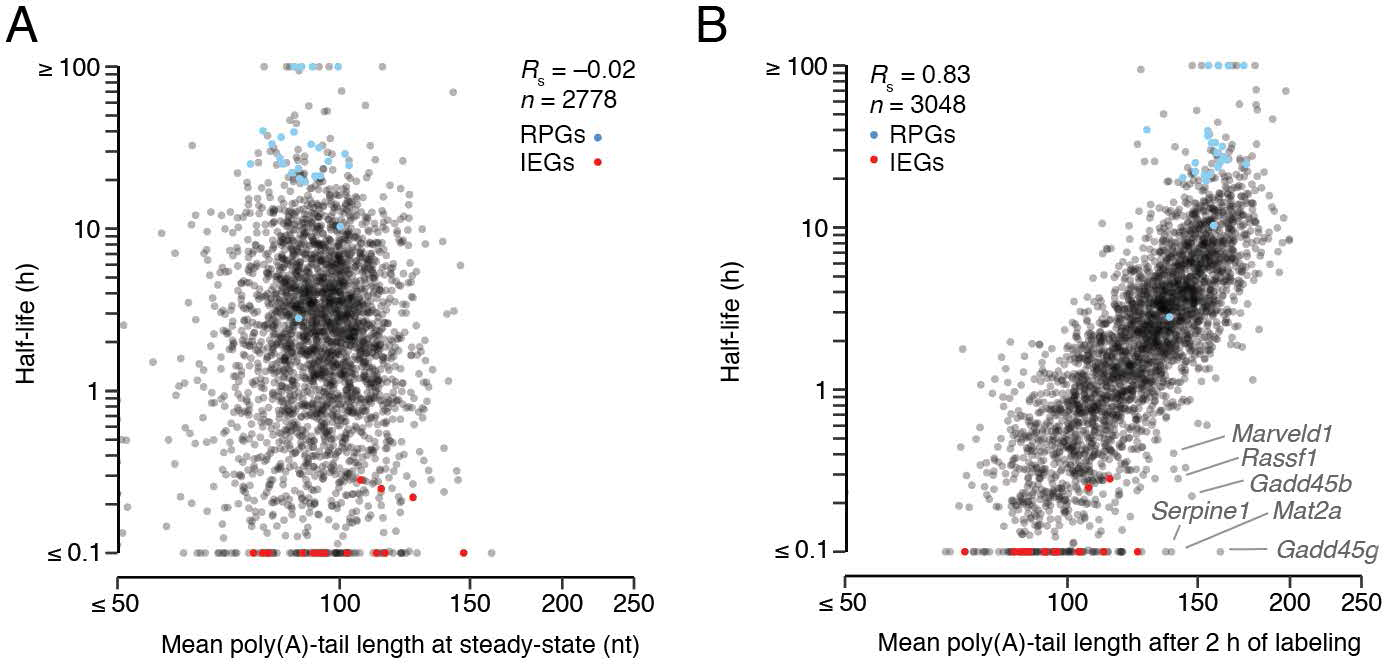
Correspondence Between mRNA Half-life and Deadenylation Rate. (A) Relationship between half-life and mean steady-state tail length of mRNAs in 3T3 cells. For mRNAs of each gene, standard PAL-seq data were used to determine the length distribution of tails ≥ 50 nt, and data generated from a protocol that used single-stranded ligation to the mRNA 3′ termini (rather than a splinted ligation to the tail) were used to determine both the length distribution of tails < 50 nt and the fraction of tails < 50 nt. Compared to the tail-length distribution generated by only standard PAL-seq data, this composite distribution better accounted for very short and highly modified tails. Nonetheless, using the standard PAL-seq data without this adjustment produced a similar result (Figure S3G). Results for mRNAs of ribosomal protein genes (RPGs) and immediate early genes (IEGs) (Tullai et al., 2007) are indicated (blue and red, respectively). (B) Relationship between mRNA half-life and mean tail length of metabolically labeled mRNAs isolated after 2 h of labeling. Otherwise as in (A). See also Figures S3A–D, S3G.

A very different picture emerged when considering pre-steady-state tail-length measurements. After 2 h of labeling, half-life strongly corresponded to mean tail length (Figure 2B; *R*_s_ = 0.83). At this labeling interval, IEG mRNAs and other short-lived mRNAs had the shortest mean tail lengths, RPG mRNAs and other long-lived mRNAs had the longest mean tail lengths, and other mRNAs had mean tail lengths falling somewhere in between. The simplest explanation for this result is that the deadenylation rate dictates the stability of most mRNAs, and mean tail length at 2 h provides a proxy for deadenylation rate. Thus, slow deadenylation of RPG mRNAs and other long-lived mRNAs explains both why they have longer tails after 2 h of labeling and why they have such long half-lives, and rapid deadenylation of IEG mRNAs and other short-lived mRNAs explains why they have shorter tails after 2 h of labeling and why they have such short half-lives.

Several notable outliers had half-lives that were shorter than expected from their mean tail lengths, suggesting that their degradation and deadenylation rates were incongruous. *Rassf1*, *Mat2a*, *Serpine1*, and two *Gadd45* paralogs are known or suspected substrates for either nonsense-mediated decay (NMD) or other pathways that recruit UPF1 (Forrest et al., 2004; Tani et al., 2012; Park and Maquat, 2013; Bresson et al., 2015; Nelson et al., 2016). Another outlier, the *Marveld1* mRNA, has not yet been reported to interact with UPF1, but its protein product does interact with UPF1 in human cells and regulates UPF1 activity (Hu et al., 2013). Association with UPF1 can trigger endonucleolytic cleavage of mammalian mRNAs, which would decouple the rates of decay and deadenylation (Muhlemann and Lykke-Andersen, 2010), disrupting the relationship between half-life and tail length at intermediate labeling intervals. Nonetheless, the most notable feature of the outliers was their scarcity; the striking overall correspondence observed between half-life and mean tail lengths after 2 h of labeling implied that for the vast majority of endogenous mRNA molecules of mouse fibroblasts, the rate of mRNA deadenylation largely determines the rate of degradation.

### Initial Tail Lengths of Cytoplasmic mRNAs

Analysis of tail-length distributions for individual genes and the changes in these distributions over increased labeling intervals supported and extended the conclusions drawn from global analyses of abundances and mean tail lengths. This analysis confirmed that tail-length dynamics of mRNAs with short half-lives (e.g., *Metrnl*) substantially differed from those of mRNAs with longer half-lives (e.g., *Lsm1* and *Eef2*), with the short-lived mRNAs reaching their steady-state abundance and tail-length distribution much more rapidly (Figure 3). The stacked pattern of the distributions observed over increasing time intervals also illustrated that the longest-tailed mRNAs observed at steady state were essentially all recently transcribed, whereas the shortest-tailed mRNAs were mostly the oldest mRNAs (Figure 3).

**Figure 3.**
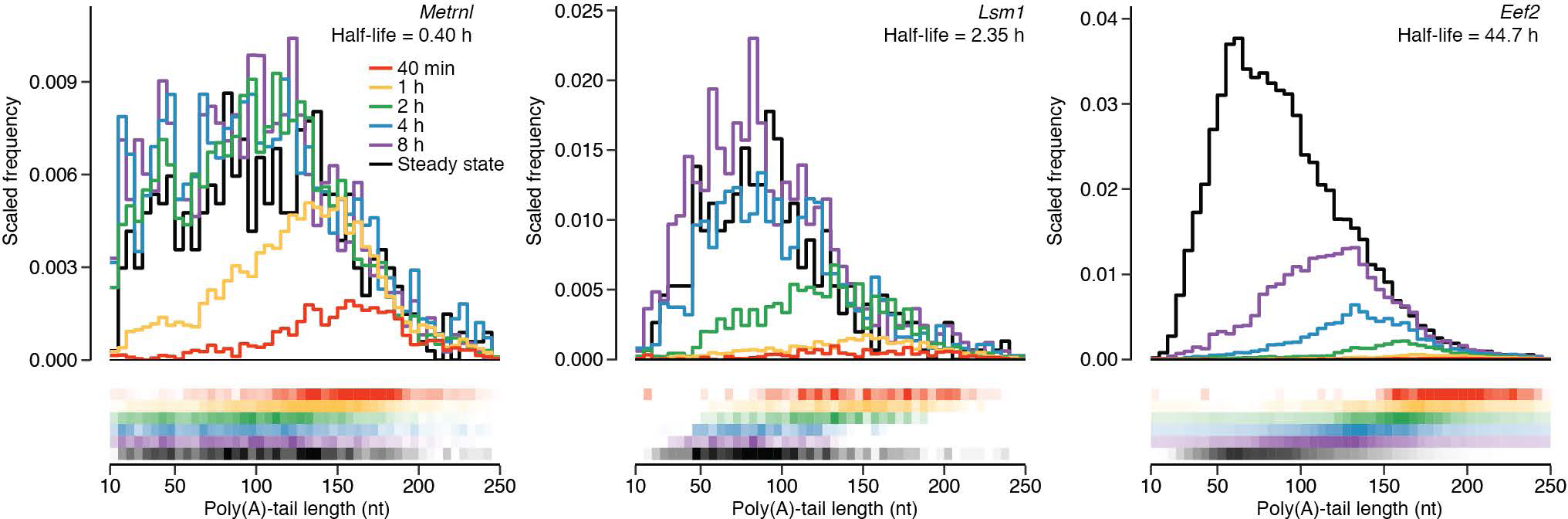
Tail-Length Dynamics of mRNAs with Different Half-Lives. Tail-length distributions for mRNAs from individual genes. For each time interval (key), the distribution is scaled to the abundance of labeled RNA in the sample (top), and the distribution is represented as a heatmap (bottom), with the range of coloration corresponding to the 5–95 percentile of the histogram density. Each bin is 5 nt. Bins for tails < 10 nt are not shown because the splinted ligation to the tail used in the standard PAL-seq protocol depletes measurements for tails < 8 nt. Bins for tails ≥ 250 nt are also now shown. See also Figures S3F, S3H–J.

Our tail-length data from short labeling periods provided the opportunity to examine the initial tail lengths of mRNAs soon after they entered the cytoplasm. The calculated 27–36 min delay in the appearance of labeled cytoplasmic mRNAs implied that most mRNAs isolated after 40 min of labeling were subject to cytoplasmic deadenylation for < 13 min. Thus, for all but the most rapidly deadenylated mRNAs, the tail lengths observed after 40 min of labeling should have approximated the tail lengths of mRNAs that first entered the cytoplasm.

Without data to the contrary, previous studies of tail-length dynamics have assumed that initial cytoplasmic tail lengths observed for mRNAs of one gene also apply to the mRNAs of all other genes. However, we observed substantial intergenic variation for average tail lengths at the shortest labeling period (Figure 1C, Figure 3, and Figure S3F), with the spread of the 5^th^ to 95^th^ percentile values at least that of steady state (112.2 ± 4.7 to 194.7 ± 6.0 nt for the 40 min samples and 84.8 ± 1.3 to 124.6 ± 2.1 nt for the steady-state samples, respectively, values ± s.d.), which suggested that mRNAs from different genes exit the nucleus with tails of quite different lengths. To examine whether deadenylation occurring soon after nucleocytoplasmic export might have influenced this result, we focused on mRNAs with half-lives > 8 h. On average, mean tail lengths for these genes exhibited less than 4% change when comparing the 40 min and 1 h time intervals, implying that they also underwent little cytoplasmic deadenylation during the first 40 min of labeling. Average tail lengths observed at 40 min for mRNAs from these genes spanned a broad range, exceeding that observed at steady state (spread of the 5^th^ to 95^th^ percentile values 128.3 ± 5.2 to 242.1 ± 16.1 nt for the 40 min samples and 81.0 ± 1.0 to 119.4 ± 1.4 nt for the steady-state samples, respectively, values ± s.d.), although these tail-length values observed at 40 min had little correspondence with those observed at steady state (*R*_s_ = 0.12).

When comparing mRNAs from the same gene, tail-length distributions were also quite broad for the newly exported mRNAs, as illustrated for mRNAs from three genes (Figure 3), and further demonstrated by the mean coefficient of variation (c.v.) of 0.41 for mRNAs of all measured genes (Figure S3H), compared to a c.v. of 0.20 for the 160 nt standard in the 40 min sample. These c.v. values were reproducible between biological replicates and had little correspondence with mRNA half-life (Figure S3I–J). Although we cannot rule out the formal possibility that mRNA tails undergo exceedingly rapid and variable transient deadenylation immediately upon nuclear export, we interpret our results at short labeling periods to indicate that mRNAs exit the nucleus with considerable but reproducible intergenic and intragenic tail-length variability.

### A Quantitative Model of mRNA Deadenylation and Decay

Our ability to isolate mRNAs of different age ranges for each gene and analyze their abundance and tail lengths (Figure 3) provided the unique opportunity to calculate the deadenylation rates and other metabolic rates and parameters for these mRNAs, thereby expanding the number of metabolically characterized mammalian mRNAs far beyond the four (*Mt1, Fos*, *Hbb*, and *IL8*) that have been examined using single-gene measurements (Mercer and Wake, 1985; Wilson and Treisman, 1988; Shyu et al., 1991; Gowrishankar et al., 2005). In contrast to mRNA half-lives, which are fit directly to the approach to steady state, other rates are not as simple to calculate. For each gene, the number of mRNA molecules with a given tail length is a function of 1) the rate of mRNA entering the cytoplasm, a function of the rates of transcription, processing, and nucleocytoplasmic export, 2) the tail-length distribution of mRNA entering the cytoplasm, 3) the deadenylation rate, 4) the tail length below which the mRNA is no longer protected from decapping, and 5) the decapping rate of short-tailed mRNAs (with decapping assumed to trigger rapid decay of the mRNA body). Therefore, we developed a mathematical model to determine, for mRNAs of thousands of genes, values for each of these parameters.

Our model was based on a system of differential equations that describe the rates of change of abundance of mRNA intermediates (Figure 4A and Table S2), an approach resembling that used to model metabolism of RNAs from single-gene reporters (Cao and Parker, 2001; Jia et al., 2011). For each gene, transcription, nuclear processing, and export (hereafter abbreviated as “production”) generates, with rate constant *k*_0_, a distribution of initial poly(A)-tail lengths. Over time, deadenylation shortens the tail, one nucleotide at a time, with rate constant *k*_1_. Decapping, with rate constant *k*_2_, can occur alongside deadenylation and monotonically increases as the poly(A)-tails get shorter. Because the body of each decapped mRNA is rapidly degraded (instantly in our model), decapping reduces the abundance of mRNAs in the pool. Note that although this model parameter is named “decapping” based on the idea that mRNA decay proceeds primarily through decapping and subsequent 5′-to-3′ decay, decay of a short-tailed mRNA body by other mechanisms would also be consistent with our model.

**Figure 4.**
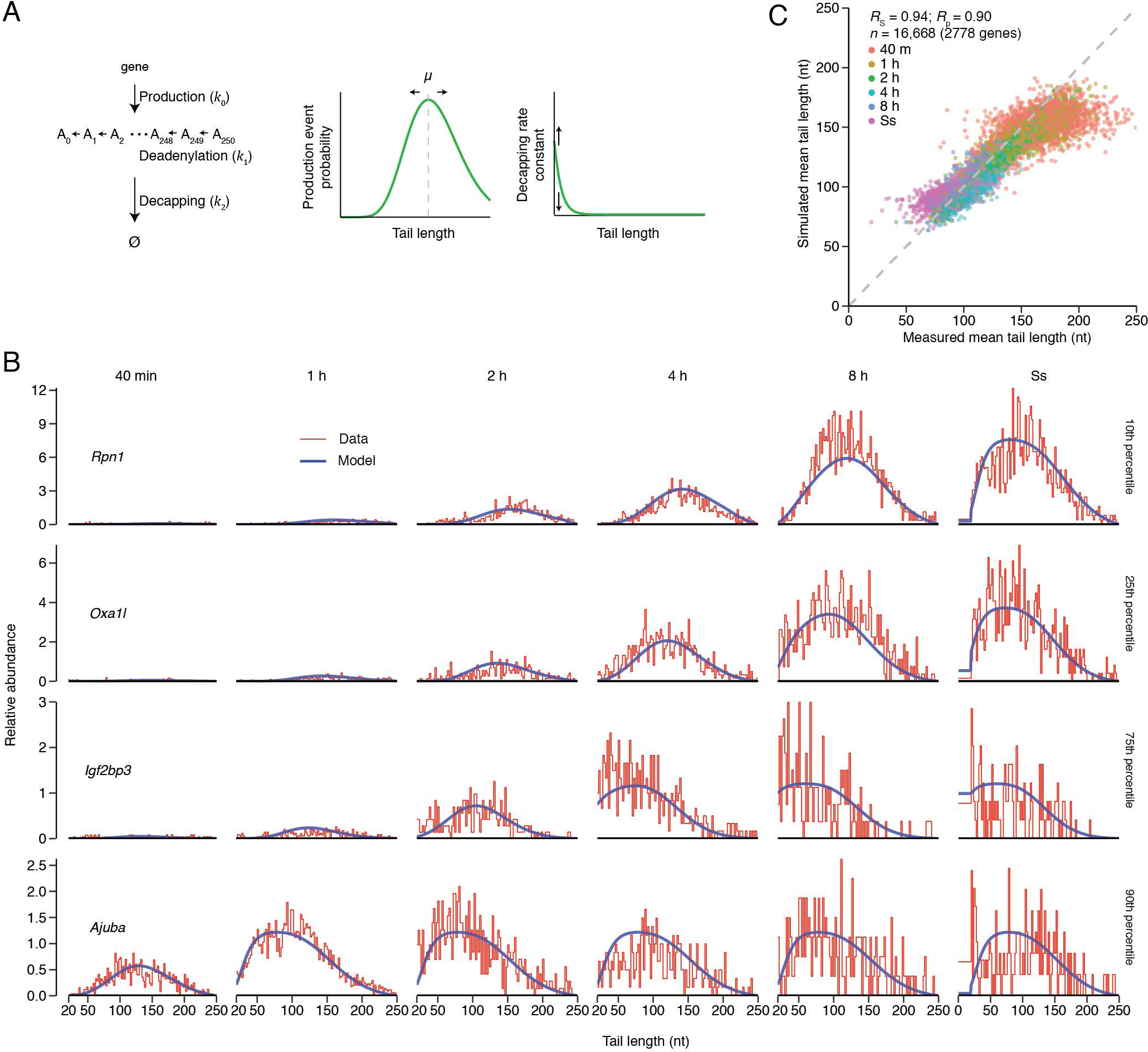
Computational Model of mRNA Deadenylation and Decay Dynamics. (A) Schematic of the computational model. Poly(A)-tail lengths of mRNAs are represented by A*_n_*, where *n* is the length of the poly(A) tail. *k*_0_*, k*_1_, and *k*_2_ are terms for mRNA production, deadenylation, and decapping, respectively, and Ø represents the loss of the mRNA molecule. The curves (right) indicate the distributions used to model probabilities of production and decapping as functions of tail length. They are schematized using the globally fitted parameters (*v_p_, m_d_*, and *v_d_*) that defined each distribution (Table S2). The parameter *m_p_* controls the mean (*µ*) of the negative binomial distribution (left curve), whereas the decapping rate constant, *β*, scales the decapping distribution (right curve) (Table S2). (B) Correspondence between the fit of the model and the experimental data. Results for mRNAs of these four genes are shown as representative examples because their fits fell closest to the 10^th^, 25^th^, 75^th^, and 90^th^ percentiles of the distribution of *R*^2^ values for all genes that passed expression cutoffs in the PAL-seq datasets (Figure S4F, *n* = 2778). For each time interval, the blue line shows the fit to the model, and the red line shows the distribution of observed tail-length species, plotted in 2 nt bins and scaled to standards as in Figure 1B. (C) Correspondence between mean tail lengths generated from the model simulation and tail lengths measured in the metabolic labeling experiment. Shown for each gene are mean tail lengths for mRNAs at each time interval (key) from the simulation plotted as a function of the values observed experimentally. The discrepancy observed for some mRNAs at early time intervals was attributable to low signal for long-lived mRNAs at early times. The dashed line indicates *y* = *x*. See also Figures S4, S5A–B, and Tables S1–S2.

For individual mRNAs generated from the same gene, the production terms varied according to a negative binomial distribution—a distribution routinely used to model the probability of a failure after a series of successes (in our case, creating an mRNA of tail length *n* + 1 after successfully creating an mRNA of tail length *n*) (Figure 4A and Table S2). The decapping rate constant followed a logistic function, which accelerated decapping as tails shortened. The two parameters of this function (*m_d_* and *v_d_*) were fit as global constants, while the scaling parameter (*β*) was fit to each gene (Table S2). Solving the differential equations of the model estimated both the tail-length distribution and the mRNA abundance at each time interval for mRNAs from each gene.

Before arriving at the final version of the model (Figure 4A), we considered alternative models with varying levels of complexity. For example, building on the proposal that most mRNAs are substrates for both the Pan2/Pan3 and Ccr4/Not deadenylase complexes, with Pan2/Pan3 acting on tails > 110 nt and Ccr4/Not acting on shorter tails (Yamashita et al., 2005), we tested the performance of a model with two deadenylation rate constants, in which the transition between the two occurred at a tail length of 110 nt (Figure S4A). This model yielded residuals that were only marginally improved (Figure S4B), and for each mRNA the two deadenylation rates resembled each other (Figure S4C). A model in which the transition between the deadenylation rates occurred at 150 nt (Yi et al., 2018) yielded similar results (Figure S4D–E). These results indicated that, for endogenous mRNAs in 3T3 cells, either a single deadenylase complex dominates—as recently proposed for mRNAs with tail lengths ≤ 150 (Yi et al., 2018)—or both complexes act with indistinguishable kinetics. Thus, we chose not to implement a more complex model with two deadenylation rate constants.

Fitting the final version of the model to the tail-length and abundance measurements for mRNAs from thousands of genes yielded average initial tail lengths and rate constants for production, deadenylation, and decapping for each of these mRNAs (Table S2). The correspondence between the output of the model and the experimental measurements is illustrated for genes selected to represent different quantiles of fit based on the distribution of *R*^2^ values (Figure 4B and Figure S4F). Mean tail-length values generated by the model corresponded well to measured values (Figure 4C, *R*_s_ = 0.94, *R*_p_ = 0.90). Moreover, values fit for starting tail length, production, deadenylation, and decapping were reproducible between biological replicates and robust to parameter initialization as well as multinomial sampling (bootstrap analysis) (Figure S4G–J).

### The Dynamics of Cytoplasmic mRNA Metabolism

Of the six yeast mRNAs and four mammalian mRNAs that have been metabolically characterized, the data for four yeast mRNAs and two mammalian mRNAs are of sufficient resolution to derive deadenylation rates. The two mammalian mRNAs, *Fos* and *Mt1*, have deadenylation rate constants that differ by 60-fold (20 and 0.33 nt/min, respectively) (Mercer and Wake, 1985; Shyu et al., 1991). Our analysis, which metabolically characterized 2778 mammalian mRNAs, greatly expanded the set of mRNAs with measured deadenylation rates and showed that deadenylation rate constants of mammalian mRNAs can differ by > 1000-fold—as fast as > 30 nt/min and as slow as 0.03 nt/min (Figure 5A). Concordant with our direct analysis of the primary data, which revealed a strong correspondence between mRNA half-life and pre-steady-state mean tail length, thereby implying that most mRNAs degrade through a mechanism involving tail shortening (Figure 1F), mRNA half-lives corresponded strongly to deadenylation rate constants fit to our model (*R*_s_ = –0.95, Figure S5A).

**Figure 5.**
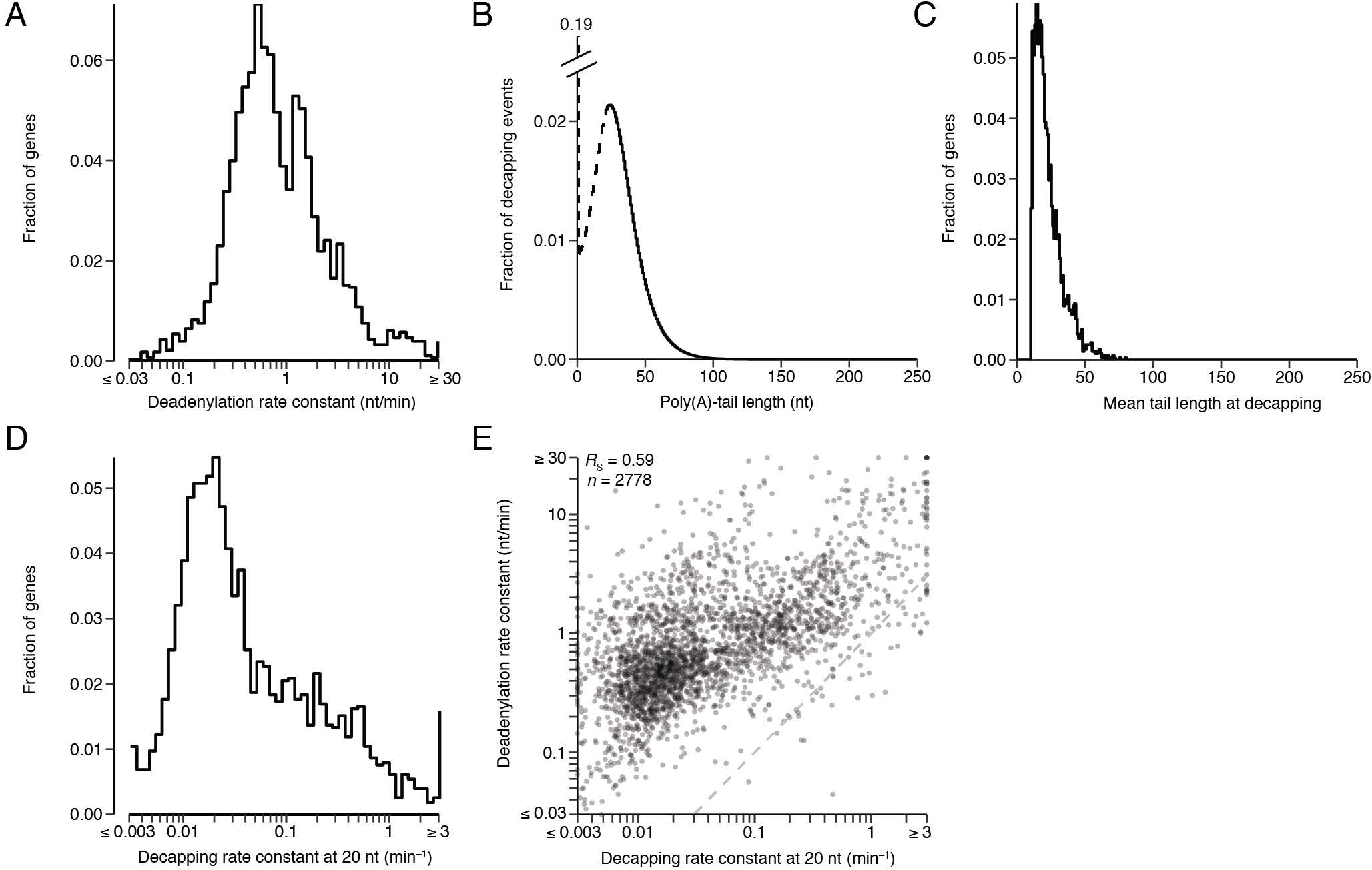
Dynamics of Cytoplasmic mRNA Metabolism. (A) Distribution of deadenylation rate constants (*k*_1_ values), as determined by fitting the model to data for mRNAs from each gene (n = 2778). (B) Tail lengths at which mRNAs are decapped, as inferred by the model. The model rate constants were used to simulate a steady-state tail-length distribution for each gene. The abundance of each mRNA intermediate was then multiplied by the decapping rate constant *k*_2_ to yield a distribution of decapping events over all tail lengths. Plotted is the combined distribution for all mRNA molecules of all 2778 genes. Results were indistinguishable when the distribution from each gene was weighted equally. Values for tails < 20 nt are shown as a dashed line because the model fit steady-state tail lengths < 20 nt as an average of the total abundance of tails in this region, and thus did not provide single-nucleotide resolution for decapping rates of these species. (C) Mean tail lengths at which mRNAs from each gene (n = 2778) were decapped, as inferred by the model. Otherwise, as in (B). (D) Distribution of decapping rate constants (*k*_2_ values) for mRNAs with 20 nt tail lengths, as determined by fitting the model to data for mRNAs from each gene (n = 2778). (E) Correlation between the deadenylation rate constant (*k*_1_) and the decapping rate constant (*k*_2_) at a tail length of 20 nt. The dashed line indicates *y* = *x*. See also Figure S4.

Our model and its fitted parameters allowed us to compute the decapping rates for all measured genes at all tail lengths and thereby infer the tail lengths at which mRNAs were decapped and degraded (Figure 5B). This analysis indicated that nearly all decapping occurred after the tail lengths fell below 100 nt, which agreed with previous analyses of reporter genes (Yamashita et al., 2005). Decapping greatly accelerated as tail lengths fell below 50 nt (with > 92% of mRNAs decapped below this length), a length less than the 54 nt footprint of two cytoplasmic poly(A)-binding protein (PABPC) molecules (Baer and Kornberg, 1983; Yi et al., 2018). However, most mRNA molecules (> 55%) were not decapped until their tail lengths fell below 25 nt, a length less than the 27 nt footprint of a single PABPC molecule (Figure 5B).

When analyzing the mean tail length of decapping for mRNAs of each gene, the results generally concurred with those observed for all mRNA combined, with mRNAs from most genes decapped at short mean tail lengths (Figure 5C, > 97% decapped at mean tail length < 50 nt and > 69% decapped at mean tail length < 25 nt). As expected, most mRNAs previously found to have discordant deadenylation and decay rates (Figure 1F) were also outliers in this analysis, with *H2afx*, *Mat2a*, *Gadd45b*, and *Marveld1*, degrading at a mean tail length of 75, 70, 62, and 59 nt, respectively. The estimates of mean decapping tail lengths together with initial tail lengths and deadenylation rate constants enabled estimates of the time required to reach the mean tail length of decapping, which corresponded to lifetime slightly better than did the deadenylation rate constants on their own to half-life (Figure S5A–B, *R*_s_ = –0.96 and –0.95, respectively.)

Once tails reached a short length, the decapping rate constants of short-tailed mRNAs varied widely, with short-tailed mRNAs from some genes undergoing decapping at rate constants > 1000-fold greater than those of short-tailed mRNAs from other genes (Figure 5D). *Fos*, a rapidly deadenylated mRNA, is degraded much faster upon reaching a short tail length than is *Hbb*, a less rapidly deadenylated mRNA (Shyu et al., 1991). If general for other mRNAs, a more rapid degradation of short-tailed mRNAs that had been more rapidly deadenylated would help prevent buildup of short-tailed isoforms of rapidly deadenylated mRNAs. However, such buildup sometimes does occur, as observed in *Drosophila* cells for three mRNAs characterized during heat shock (Dellavalle et al., 1994; Bönisch et al., 2007) and in mammalian cells for *Csf2* (Chen et al., 1995; Carballo et al., 2000), raising the question of the extent to which decay rates of short-tailed mRNAs are coupled to their deadenylation rates. To answer this question, we examined the relationship between rate constants for deadenylation and those for decay of short-tailed mRNAs (the latter calculated for mRNAs with 20 nt tails). As reported for *Fos*, more rapidly deadenylated mRNAs tended to be degraded more rapidly upon reaching short tail lengths (Figure 5E, *R*_s_ = 0.59).

### A Modest Buildup of Short-Tailed Isoforms of Short-Lived mRNAs

Having found this more rapid clearing of mRNAs that had been more rapidly deadenylated, we investigated whether this phenomenon was able to prevent a large build-up of short-tailed isoforms of rapidly deadenylated mRNAs. For this investigation, we analyzed the steady-state dataset that incorporated results of PAL-seq implemented with direct ligation to mRNA 3′ termini, which better detected very short or highly modified tails. Despite the rapid decay of short-tailed mRNAs that had been more rapidly deadenylated, less stable mRNAs generally did have a somewhat higher fraction of short-tailed transcripts (Figure 6A and Figure S5C, *R*_s_ = –0.56). Nonetheless, the buildup of short-tailed isoforms of these unstable RNAs usually failed to exceeded 30% of all transcripts (Figure 6A).

**Figure 6.**
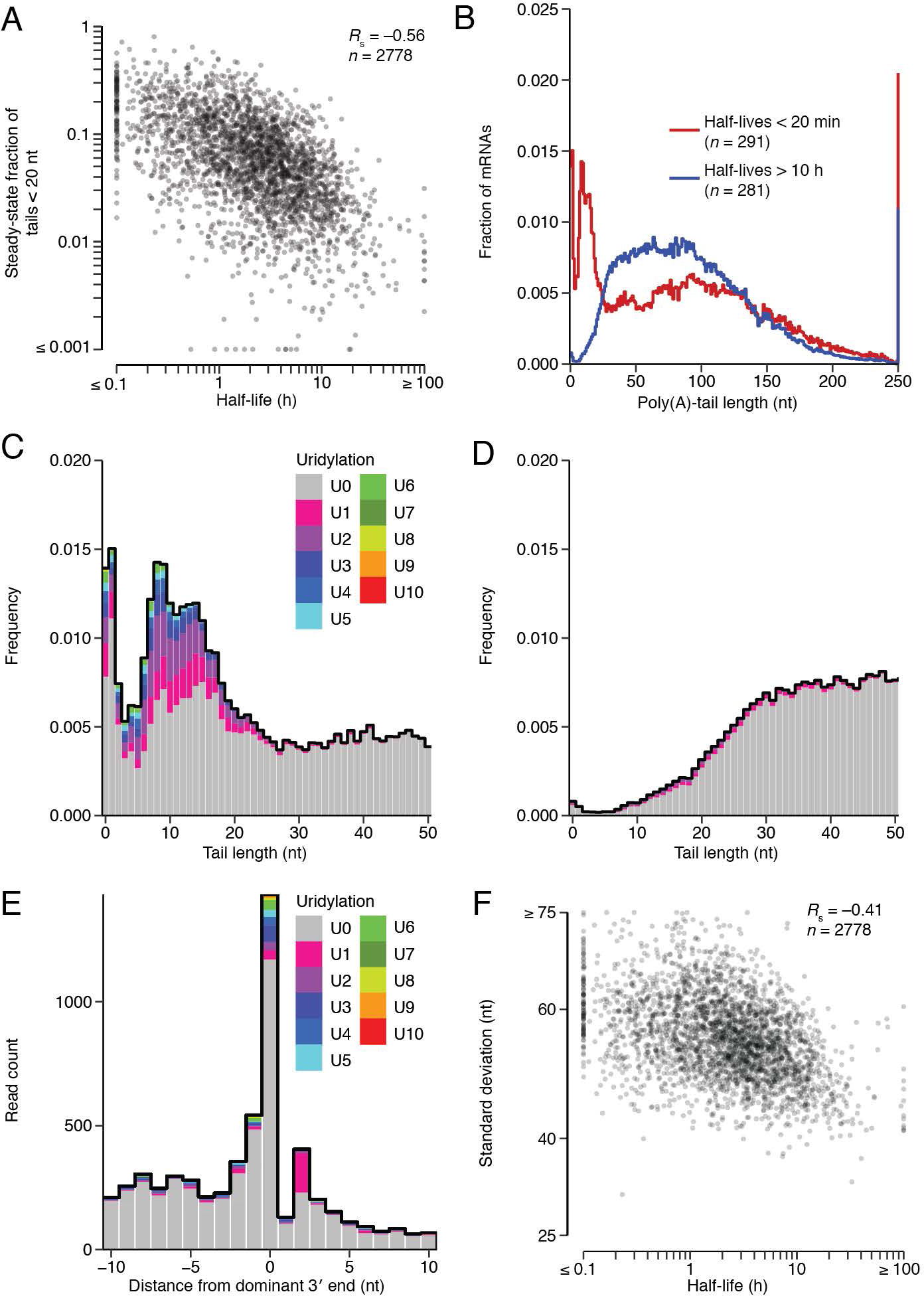
A Modest Buildup of Short-Tailed Isoforms of Short-Lived mRNAs. (A) Relationship between the steady-state fraction of tails < 20 nt and mRNA half-life. For mRNAs of each gene, the fraction of tails < 20 nt was calculated from a composite distribution generated as in Figure 2A, which accounted for very short and highly modified tails. (B) Metatranscript distributions of steady-state tail lengths of short- and long-lived mRNAs (red and blue, respectively), with mRNAs from each gene contributing density according to their abundance. Results were almost identical when mRNAs were weighted such that each gene contributed equally. This analysis used the composite distributions as in (A). (C) Uridylation of short-lived mRNAs with short poly(A) tails. For mRNAs with half-lives < 20 min, the fraction of molecules with the indicated poly(A)-tail length at steady-state is plotted, indicating for each tail length the proportion of tails appended with 0 through 10 U nucleotides (key). For mRNAs with poly(A)-tail length of 0, U residues were counted only if they could not have been genomically encoded. As poly(A) tails approached 20 nt the ability to map reads with ≥ 3 terminal U residues diminished, but the ability to map reads with 1–2 terminal U residues was retained for poly(A) tails of each length. (D) Uridylation of long-lived mRNAs (half-lives > 10 h) with short poly(A) tails. Otherwise as in (C). (E) Distribution of tailless tags (regardless of half-life) as a function of their distance from the annotated 3′ end of the UTR. Tags with a terminal A (or with a terminal A followed by one or more untemplated U) were excluded, even if the A might have been genomically encoded. The proportion of tails appended with 0 through 10 U nucleotides is shown (key). (F) Relationship between the standard deviation of steady-state tail length and mRNA half-life. Otherwise as in (A). See also Figure S5C–J.

This preferential buildup of short-tailed isoforms of unstable RNAs was clearly visualized in a meta-transcript analysis of the tail-length distribution at steady state. Short-lived mRNAs (half-lives < 20 min) had two peaks of short-tailed isoforms, a major peak centering at 7–15 nt and a minor peak at 0–2 nt, whereas long-lived mRNAs (half-lives > 10 h) were depleted of tails of < 20 nt (Figure 6B). Closer inspection of these two peaks revealed that these short-tailed isoforms of short-lived mRNAs were dramatically enriched in mono- and oligouridylated termini (Figure 6C, D and Figure S5D), consistent with studies showing that uridylation occurs preferentially on shorter tails and helps to destabilize mRNAs (Kwak and Wickens, 2007; Rissland et al., 2007; Rissland and Norbury, 2009; Chang et al., 2014; Lim et al., 2014), and further suggesting that uridylation preferentially occurs on short-lived mRNAs.

The observation of a 0–1 nt peak in the steady-state tail-length distribution prompted examination of fully deadenylated isoforms of mRNAs that were initially polyadenylated. Molecules without tails were often also missing the last few nucleotides of the 3′ UTR (Figures 6E and Figure S5E), suggesting that after removing the tail, the deadenylation machinery (or some other 3′-to-5′ exonuclease) usually proceeds several nucleotides into the mRNA body. Analysis of mRNAs with tails indicated that, with few exceptions, the last nucleotide of the 3′ UTR was consistently defined (Figure S5F–H), which supported the idea that the missing nucleotides of tailless molecules had not been lost during the process of cleavage and polyadenylation. Analysis of the final dinucleotides of tailless tags revealed no consistent pattern after accounting for the genomic background, suggesting that other factors, such as proteins or more distal nucleotide composition, influence the position at which the exonuclease stops.

Despite their presence, the two peaks of short-tailed isoforms did not dominate the distribution, as most short-lived mRNAs (70%) had tails exceeding 30 nt (Figure 6B). Indeed, compared to long-lived mRNAs, these short-lived mRNAs also had modest enrichment for very long tails (> 175 nt) (Figure 6B and Figure S5I–J), perhaps due to an initial lag in assembling deadenylation machinery as mRNAs enter the cytoplasm, which would the cause a relatively larger fraction of short-lived mRNAs to exist in the cytoplasm prior to an initial encounter with a deadenylase. The increased fractions of both short-tail and long-tail isoforms for short-lived mRNAs led to broader overall tail-length distributions (Figure 5B) with increased standard deviations in tail length (Figure 6F, *R*_s_ = –0.41). Moreover, the increased fractions of shorter and longer isoforms offset each other when calculating mean tail length, leading to similar mean tail lengths for the short- and long-lived mRNAs (Figure S5K, median mean tail lengths = 89 and 92 nt for short- and long-lived mRNAs, resepectively), which contributed to the lack of correlation between half-life and mean tail length at steady state (Figure 2A). Most importantly, the low magnitude of the buildup supported our conclusion that for most mRNAs the steps of deadenylation and subsequent decay are kinetically coupled: short-tailed mRNAs that had previously undergone more rapid deadenylation are more rapidly decayed. This coupling prevents a large buildup of short-tailed isoforms of rapidly deadenylated RNAs, thereby enabling the large range in deadenylation rate constants to impart a similarly large range in mRNA stabilities.

### Deadenylation and Decay Dynamics of Populations of Synchronous mRNAs

Our continuous-labeling experiments were designed to measure dynamics of mRNA metabolism in an unperturbed cellular environment. However, this framework required deadenylation and decapping parameters to be inferred as mRNAs from each gene approached their steady-state expression levels and tail lengths, with their populations becoming progressively less synchronous, causing the signal for the end behavior of mRNAs to be diluted. For orthogonal measurements of these parameters, we performed a pulse-chase–like experiment that more closely resembled previous studies with single-gene reporters, in that it monitored synchronous populations of mRNAs from each gene. After a 1 h pulse of 5EU, 3T3 cells were treated with actinomycin D (actD) to block transcription, and the abundances and poly(A)-tail lengths of the mRNAs produced during the 5EU-labeling period were measured over the next 15 h, thereby revealing the behavior of synchronized mRNA populations as they age (Figure 7A).

**Figure 7.**
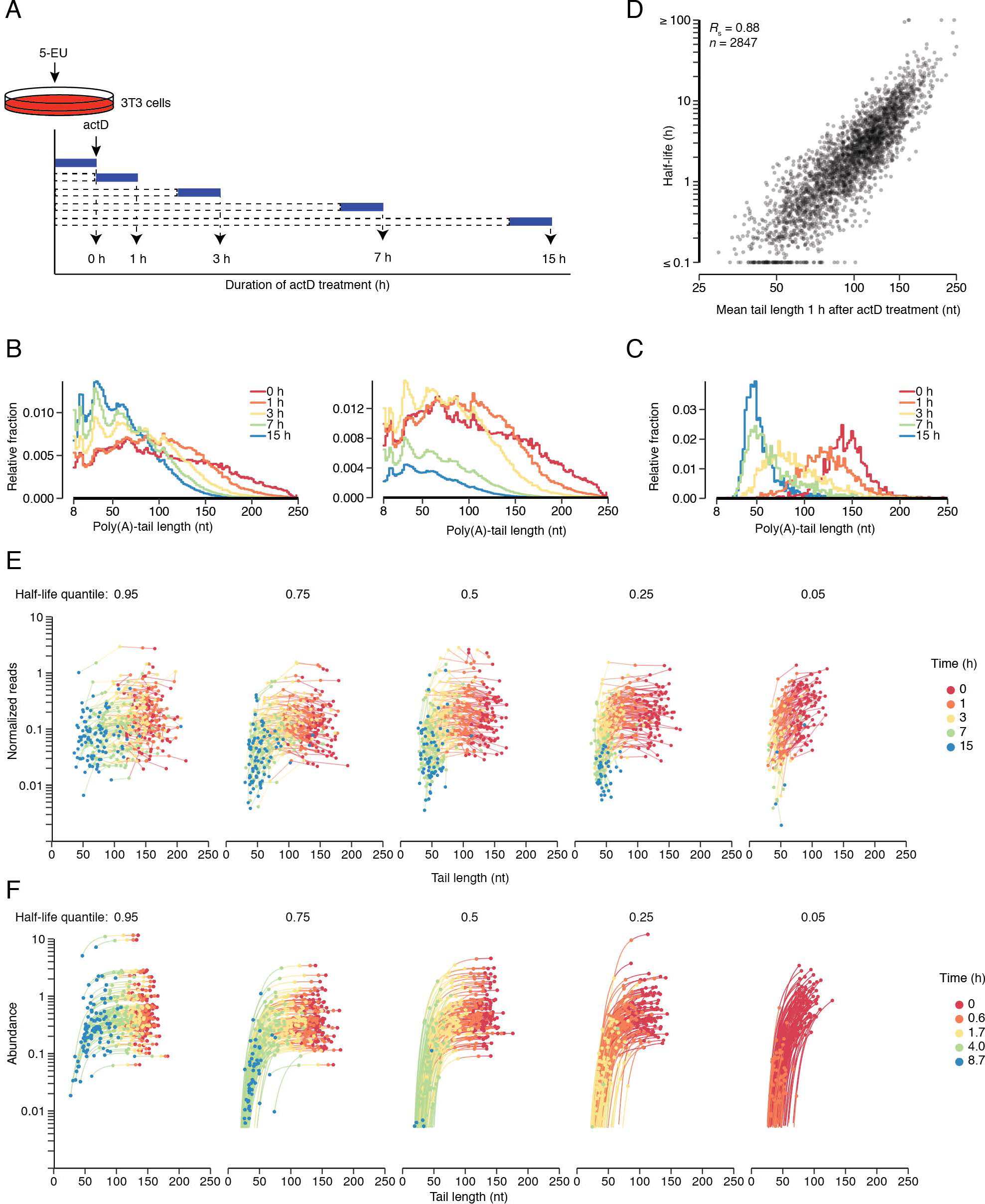
Deadenylation and Decay Dynamics of Synchronous mRNA Populations. (A) Schematic of 5EU metabolic-labeling and actD treatments used to analyze synchronized cellular mRNAs. Cells from cell line 2 were treated for 1 h with 5EU, then treated with actD continuously over a time course spanning 15 h. (B) Tail-length distributions of labeled mRNA molecules observed at the indicated times after stopping transcription (key). Left: Distributions were normalized to all have the same area. Right: Distributions were scaled to the abundance of labeled RNA in each sample and then normalized such that the 0 h time interval had an area of 1. Each bin is 2 nt; results for the bins with tail lengths < 8 nt and ≥ 250 nt are not shown. At 0 h, 7% of the tails were still ≥ 250 nt, which helps explain why the density for the remainder of the tails fell below that observed at 1 h. (C) Distributions of mean poly(A)-tail lengths for labeled mRNAs of each gene after the indicated duration of transcriptional shutoff. Values for all mRNAs that passed the cutoffs for tail-length measurement at all time points were included (n = 2155). Each bin is 2 nt. (D) Relationship between half-life and mean tail length of labeled mRNAs from each gene after 1 h of actD treatment. (E) Labeled mRNA abundance as a function of mean tail length over time. Results are shown for mRNAs grouped by half-life quantiles (95%, 75%, 50%, 25%, and 5%, left to right, with mRNAs in the 5% bin having shortest half-lives). Each half-life bin contains 100 genes. mRNA abundance was determined from paired RNA-seq data. Each line connects values for mRNA from a single gene. (F) Simulation of mRNA abundance as a function of mean tail length over time. For each gene in (E), model parameters fit from the continuous-labeling experiment were used to simulate the initial production of mRNA and its mean tail length from each gene, as well as the fates of these mRNAs and mean tail lengths after production rates were set to 0. Results are plotted as in (E), but using a shorter time course (key) to accommodate the faster dynamics observed without actD. See also Figure S3E.

As expected, the tail lengths of labeled mRNAs progressively decreased after transcriptional inhibition, with median tail lengths shortening from 123 to 51 nt over the course of the experiment (Figure 7B). Examination of mean tail lengths of mRNAs from each gene revealed a similar trend (Figure 7C). At later time points the mean tail-length distributions peaked between 45–50 nt (Figure 7C), far below the 100–105 nt mode of the steady-state distribution, which included mRNAs of all ages (Figure 1C).

The actD treatment had some side effects. At later time points, a ∼30 nt periodicity emerged in the single-molecule tail-length distributions (Figure 7B). Although such phasing of tail lengths, with a period resembling the size of a PABPC footprint, has been observed in mammalian cells following CCR4 knockdown (Yi et al., 2018) and in *C. elegans* (Lima et al., 2017), only subtle phasing was observed in unperturbed mammalian cells (Figure 6B). This more prominent periodicity observed after prolonged actD treatment was presumably the result of more dense packing of PABPC on poly(A) tails in the context of a diminishing mRNA pool. A second side effect of actD treatment concerned mRNA half-lives, which increased from a median of 2.1 h in the continuous-labeling experiment to a median of 3.8 h in the transcriptional-shutoff experiment (Figure S3E). This increase was observed even for the mRNAs with the shortest half-lives, which indicated that it occurred before actD could have influenced protein output, i.e., in less time than that required for mRNA nucleocytoplasmic export and translation. This result generalized previous observations concerning the effects of actD on reporter-mRNA stabilities (Chen et al., 1995).

Despite the side effects of actD, the rank order of mRNA half-lives determined from the transcriptional-shutoff experiment agreed well with that from the continuous-labeling experiment (Figure S3E, *R*_s_ = 0.78), indicating that the transcriptional-shutoff experiment captured key aspects of the unperturbed behavior. In addition, mRNA half-lives calculated from the continuous-labeling experiment strongly corresponded to mean tail length observed 1 h after actD treatment (Figure 7D; note that 1h after actD treatment was 2 h after 5EU labeling and thus most comparable to Figure 2B). Indeed, the strength of the correspondence between half-life and 1 h tail length (*R*_s_ = 0.88) provided compelling support for our conclusion that the vast majority of mRNAs are primarily degraded through deadenylation-linked mechanisms.

To further analyze results of the transcriptional-shutoff experiment, we grouped mRNAs into cohorts based on their half-lives and monitored the abundance and average tail length of mRNAs from individual genes at each time point (Figure 7E). Regardless of mRNA half-life, tails initially shortened with little change in abundance until mean tail lengths fell below 100 nt. The minimal degradation of long-tailed molecules disfavored the idea that preferential degradation of long-tailed mRNAs might help explain the shift in tail lengths observed with increasing time intervals in the continuous-labeling experiment. As expected based on the strong correspondence between half-life and 1 h tail length (Figure 7D), mRNAs with shorter half-lives underwent more rapid tail shortening (Figure 7E). Once mean tail lengths fell below 50 nt (implying that a substantial fraction of tails fell below 25 nt), degradation accelerated. This acceleration was more prominent for mRNAs with shorter half-lives, which confirmed our conclusion that short-tailed mRNAs that had undergone more rapid deadenylation are also more rapidly degraded (Figure 7E).

To examine how well our model predicted this behavior, we used it to predict the results of the transcriptional-shutoff experiment, using the rate constants measured earlier from the continuous-labeling experiment. When simulating a shorter time course to account for the more rapid deadenylation, decapping, and decay observed without actD, the results predicted by the model agreed well with the experimental observations (*R*_s_ = 0.93 and 0.61 for mean tail length and abundance, respectively, n = 11,273 values above the abundance threshold for 2687 mRNAs), including the precipitous decline in abundance when mean tail lengths fell below 50 nt and the faster degradation of short-tailed mRNAs that had undergone faster deadenylation (Figure 7F). The striking correspondence between the predictions of the model, which had been trained on the continuous-labeling experiment, and the observations of the transcriptional-shutoff experiment validated the results and conclusions from both experiments as well as from our analytical framework.

## Discussion

Previous mechanistic studies of mRNA turnover provide information on deadenylation and degradation dynamics for four mammalian mRNAs and some derivatives, with deadenylation rates reported for two of these four (Mercer and Wake, 1985; Wilson and Treisman, 1988; Shyu et al., 1991; Chen et al., 1995; Gowrishankar et al., 2005; Yamashita et al., 2005). Our examination of the dynamics of deadenylation and degradation for mRNAs from thousands of endogenous genes provided a more comprehensive resource for deriving the principles of cytoplasmic mRNA metabolism. Initial analyses of our data revealed unanticipated intra- and intergenic variability in initial tail lengths and indicated that almost all endogenous mRNAs are degraded primarily through deadenylation-linked mechanisms, implying that the deadenylation rate of each mRNA largely determines its half-life with surprisingly little contribution from other mechanisms, such as endonucleolytic cleavage and deadenylation-independent decapping.

Mathematical modeling of our data expanded the known range in deadenylation rate constants from 60-fold to 1000-fold and showed that the link between deadenylation rate and decay generally operates at two levels. First, mRNAs with faster deadenylation rate constants more rapidly reach the short tail lengths associated with decapping and destruction of the mRNA body. With respect to the reason that short tail lengths trigger decay, our estimates of the tail lengths at which most mRNAs decay supported the prevailing view that loss of PABPC binding to the poly(A) tail enhances decay, with the idea that occupancy is somewhat lower on tails too short for cooperative binding of a PABPC dimer (tail lengths ∼50 nt) and much lower on those too short for efficient binding of a single PABPC molecule (tail lengths ∼20 nt).

This more rapid approach to short-tailed isoforms is not the whole story. mRNAs with identical 20-nt tails but from different genes can have widely different decay rate constants (1000-fold). Moreover, there is a logic to these differences—a logic conferred by the second link between deadenylation rate and decay: mRNAs that had previously undergone more rapid deadenylation decay more rapidly upon reaching short tail lengths. The coherent regulation of deadenylation and decapping rates functionally integrates mRNA turnover into a single process to ensure that mRNAs that are rapidly deadenylated are also rapidly cleared from the cell, which enables the large range in deadenylation rate constants to impart an equally large range in mRNA stabilities. With respect to mechanism, perhaps changes that occur as mRNA–protein complexes are remodeled to enhance deadenylation also recruit the decapping machinery and its coactivators. Physical connections between the Ccr4–Not deadenylase complex and the decapping complex (Haas et al., 2010; Ozgur et al., 2010; Jonas and Izaurralde, 2015) as well as the intracellular colocalization of these complexes (Parker and Sheth, 2007) presumably also help coordinate deadenylation and decapping rates.

The large differences observed for both deadenylation and decapping rate constants of mRNAs from different genes raise the question of what mRNA features might specify these differences. MicroRNAs and other factors that help recruit deadenylase complexes typically bind to sites in 3′ UTRs, implying that the presence or absence of these 3′-UTR sites helps to specify the differences (Mauxion et al., 2009; Muhlemann and Lykke-Andersen, 2010; Vlasova-St Louis and Bohjanen, 2011; Van Etten et al., 2012; Fabian et al., 2013; Leppek et al., 2013; Du et al., 2016; Bartel, 2018). However, despite the prevalence of regulatory factors that bind to 3′-UTR sites, global analyses of tandem UTR isoforms indicate that the magnitude of the differences conferred by 3′-UTR sequences in NIH 3T3 cells is relatively modest (Spies et al., 2013). Codon composition can also contribute to differences in mRNA stability, but this contribution explains only a small fraction of the variability observed for endogenous mRNAs of mammalian cells (Presnyak et al., 2015; Radhakrishnan et al., 2016; Forrest et al., 2018; Wu et al., 2019). Additional insight will be required to account more fully for the large differences in stabilities observed for different mRNAs. Our results indicate that the focus should be on sequences and processes that influence or correlate with deadenylation rates.

Our global observation that mRNAs typically degrade only after their tail lengths shorten extended to the mammalian transcriptome the notion that exponential decay is not fully appropriate for modeling mRNA degradation (Shyu et al., 1991; Cao and Parker, 2001; Trcek et al., 2011; Deneke et al., 2013). For the exponential model to be appropriate, an mRNA would need to have the same probability of decaying at any point after entering the cytoplasm. In contrast, our global analyses indicated that recently exported, long-tailed mRNAs typically undergo little if any decay, which supported the restricted-degradation model in which mRNAs are provided a discrete time window to function in the cytoplasm. During this window, the body of the mRNA is unaltered, but its age and lifespan are tracked and determined through the action of tail-length dynamics. Nonetheless, for some analyses we used the exponential model and referred to its decay parameter as ‘half-life’ when fitting abundance changes over time because in those cases a more complex model did not provide additional insight, and using mRNA half-lives is still common practice in the field. In most analyses, however, we used our mathematical model of the kinetics of deadenylation and decay to capture critical features of mRNA metabolism missed by naïve exponential decay.

Despite the utility of our mathematical model, it did not capture some finer details of mRNA metabolism. For example, it was not designed to model the burst of deadenylation that typically accompanies loss of each terminal PABPC molecule (Webster et al., 2018). However, when considering the aggregate behavior of multiple mRNAs from the same gene, these bursts become blurred, with some molecules in the burst phase and others between bursts. Accordingly, we fit a single, continuous deadenylation rate constant for the mRNAs of each gene. Likewise, we fit a single, continuous production rate constant for the mRNAs of each gene, despite the known burst behavior of transcription initiation when examined in single cells (Cai et al., 2008).

The uniform deadenylation rate constants of the model were also not suitable for capturing aspects of tail behavior that occurred as tails fell below 20 nt. For example, our analysis of steady-state data revealed buildups of isoforms of short-lived mRNAs at two tail-length ranges: 0–1 nt and 7–15 nt (Figure 6B). A model with uniform deadenylaiton rate constants can potentially explain a peak at 0 nt but not one at an intermediate tail length, such as 7–15 nt. Recognizing this limitation but still wanting to accurately account for the build-up of isoforms with tails < 20 nt observed for short-lived mRNAs, we fit the abundance of tails < 20 nt by averaging abundance over this length range and comparing this average to that predicted by the model. Several more parameters would be required to model a buildup of 7–15 nt tails, which might be warranted if further study shows that the fate of mRNAs with 7–15 nt tails differs from that of mRNAs with 0 nt tails—studies that can be contemplated now that the existence of this buildup is known.

Another aspect of mRNA metabolism remaining to be incorporated into a mathematical model is terminal uridylation. Modeling the extensive terminal uridylation of the short-tailed isoforms of short-lived mRNAs that accumulate (Figure 6C) might provide insight into the function of this modification. Previous studies report that terminal uridylation of the poly(A) tail stimulates decapping of mRNAs in *Aspergillus* and fission yeast, and of histone mRNAs in HeLa cells, and high-throughput studies link terminal uridylation more generally to mammalian mRNA stability (Mullen and Marzluff, 2008; Rissland and Norbury, 2009; Morozov et al., 2010; Chang et al., 2014; Lim et al., 2014). Based on these previous findings and the results of our analyses, we speculate that uridylation of non-histone mammalian mRNAs is preferentially deployed to short-tailed isoforms of more rapidly deadenylated transcripts to promote more rapid decapping, which helps prevent a larger buildup of these short-tailed isoforms. Regardless of the role for uridylation, the observation of these buildups of preferentially uridylated 0–1 and 7–15 nt tails for short-lived mRNAs helps explain the observations that, compared to long-lived mRNAs, short-lived RNAs have both a higher fraction of uridylated tails and a higher fraction of short-tailed isoforms that are uridylated (Chang et al., 2014; Lim et al., 2014).

A recent study observed that cytoplasmic noncanonical poly(A)-polymerases can extend tails, acting on longer-tailed mRNAs and adding mostly A residues but also sometimes generating a mixed tail in which a G or occasionally another non-A nucleotide has been incorporated (Lim et al., 2018). Because most mRNAs with these mixed tails would not be detected by PAL-seq, these mRNAs would have appeared to have been degraded in our analysis. Thus, our observation of little-to-no degradation of long-tailed mRNAs indicated that, in 3T3 cells, mRNAs with mixed tails comprised only a small fraction of the mRNA molecules at any point in time and did not impact the overall conclusions of our study.

Although our current approach does not model all aspects of mRNA metabolism, there is every reason to believe that the broad behaviors observed in these initial analyses will continue to be observed in more detailed representations of mRNA metabolism. With acquisition of suitable pre-steady-state data, the dynamics of tail-length changes in the 0–20 nt range, of terminal uridylation, and of cytoplasmic polyadenylation could be better characterized—ultimately enabling incorporation of these phenomena into a comprehensive model of mRNA metabolism. Our methods and analytical framework offer inspiration as well as a foundation for these future efforts.

## Acknowledgements

We thank J. Kwasnieski and other members of the Bartel lab for helpful discussions and the Whitehead Genome Technology Core for high-throughput sequencing. This research was supported by NIH grants GM061835 and GM118135 (D.P.B.) and an NSF Graduate Research Fellowship (T.J.E.). D.P.B. is an investigator of the Howard Hughes Medical Institute.

## Author Contributions

A.O.S., S.W.E., T.J.E., and D.P.B. conceived the project and designed the study. T.J.E., S.W.E., and A.O.S. performed the molecular experiments and analysis. T.J.E. performed the computational modeling, with input from K.S.L. and S.E.M. S.W.E., S.G., and T.J.E. adapted PAL-seq for compatibility with current Illumina technologies. K.S.L. and S.W.E. wrote the analysis pipeline for determining tail-length measurements from PAL-seq data. A.O.S., S.W.E., and T.J.E. drafted the manuscript, and T.J.E. and D.P.B. revised the manuscript with input from the other authors.

## Methods

### Cell culture

Clonal 3T3 cell lines engineered to express miR-155 (cell line 1) or miR-1 (cell line 2) upon doxycycline treatment were previously described (Eichhorn et al., 2014). Cells were grown at 37°C in 5% CO_2_ in DMEM supplemented with 10% BCS (Sigma-Aldrich) and 2 µg/mL puromycin. For metabolic-labeling time courses, cells from each line were plated onto 500 cm^2^ plates at 6.6 million cells per plate and cultured for two days such that they reached ∼70–80% confluency, at which point growth media was supplemented with 5-ethynyluridine (5EU, Jena Biosciences) (Jao and Salic, 2008) at a final concentration of 400 µM. After the desired labeling intervals cells were harvested (Figure 2A). Four plates were harvested for each 40 min time interval, three plates for each 1 h time interval, and two plates for each other time interval. A plate that had never received 5EU was harvested in parallel for each condition.

Cells were harvested at 4°C, washed twice with 50 mL ice-cold 9.5 mM PBS, pH 7.3 containing 100 µg/mL cycloheximide and then used to prepare cytoplasmically enriched lysate as described (Subtelny et al., 2014). An aliquot of cleared lysate was flash frozen for use in ribosome profiling (Eisen et al., 2019), and the rest of the lysate was added to 5 volumes of TRI reagent (Ambion) and frozen at –80°C. Samples stored in TRI reagent were thawed at room temperature, and RNA was purified according to the manufacturer’s protocol and used for RNA-seq or PAL-seq v2.

### RNA standards

Two sets of tail-length standards (set 1 and set 3, Table S3) were described previously (standard mix 2 and standard mix 1) (Subtelny et al., 2014). The other set of standards (set 2, Table S3) was prepared based on a 705 nt fragment of the *Renilla* luciferase mRNA, which was transcribed and gel purified as described (Subtelny et al., 2014) and then capped using a Vaccinia capping system (2000 µL reaction containing 500 µg RNA, 1000 U Vaccinia capping enzyme (NEB), 1X Capping Buffer (NEB), 0.1 mM S-adenosyl methionine, 0.5 mM GTP, 50 nM [α–^32^P]-GTP, 2000 U SUPERaseIn (ThermoFisher) at 37°C for 1 h), monitoring the amount of incorporated radioactivity to ensure that capping was quantitative. Following the capping reaction, the 2′,3′ cyclic phosphate at the 3′ end was removed using T4 polynucleotide kinase (Subtelny et al., 2014). The capped, dephosphorylated product was joined by splinted ligation to each of seven different poly(A)-tailed barcode oligonucleotides (Subtelny et al., 2014). These seven 3′ ligation partners included 110 and 210 nt poly(A) oligonucleotides prepared as described (Subtelny et al., 2014), and five gel-purified synthetic oligonucleotides (IDT), one with a 10 nt poly(A) tract and the other four with a 29 nt poly(A) tract followed by either A, C, G, or U. Ligation products were gel purified, mixed in desired ratios, with final ratios of the different-sized species confirmed by analysis on a denaturing polyacrylamide gel.

Short and long standards were used to monitor enrichment of 5EU-containing fragmented RNA or non-fragmented RNA, respectively. Short 5EU standards were prepared by in vitro transcription of annealed DNA oligos to produce a 30 nt and 40 nt RNA, with the latter containing a single 5EU (Table S3). In vitro transcription was performed with the MEGAscript T7 transcription kit (ThermoFisher) according to the manufacturer’s protocol, except UTP was replaced with 5-ethynyluridine-triphosphate (Jena Biosciences) when transcribing the 40 nt RNA. Long standards were prepared by in vitro transcription of sequences encoding firefly luciferase and GFP using the MEGAscript T7 transcription kit and 0.1 µM PCR product as the template. When transcribing *GFP*, a 20:1 ratio of UTP to 5-ethynyluridine-triphosphate was used. Short and long standards were gel purified and stored at –80°C. Prior to use, a portion of each standard was cap-labeled and gel purified again, which enabled measurement of the recovery of the 5EU-containing standard relative to that of the uridine-only standard.

Three 28–30 nt RNAs (Table S3) were synthesized (IDT) for use as quantification standards in RNA-seq. These standards were gel purified, and 0.1 fmol of each was added to each sample immediately prior to library preparation.

### Biotinylation of 5EU labeled RNA

The RNA-seq libraries analyzed in this study were from fragmented RNAs, size selected to match ribosome-profiling libraries (Eisen et al., 2019). For these libraries, poly(A) RNA was purified from 50 µg total RNA of the 40 min, 1, 2, and 4 h samples and 25 µg total RNA of the 8 h sample using oligo(dT) Dynabeads (ThermoFisher) according to manufacturer’s protocol. RNA was fragmented and 27–33 nt fragments were isolated as described (Subtelny et al., 2014), short standards that monitored 5EU enrichment were added, and then Cu(II) catalysis was used to biotinylate 5EU in a 20 µL reaction containing 50 mM HEPES, pH 7.5, 4 mM disulfide biotin azide (Click Chemistry Tools), 2.5 mM CuSO_4_, 2.5 mM Tris(3-hydroxypropyltriazolylmethyl)amine (THPTA, Sigma-Aldrich), and 10 mM sodium ascorbate, incubated at room temperature for 1 h. Reactions were stopped with 5 mM EDTA and then extracted with phenol–chloroform (pH 8.0). For the steady-state samples, 5 µg of RNA from the 40 min sample was poly(A) selected and fragmented, and size-selected 27–33 nt fragments were carried forward without enriching for 5EU.

For PAL-seq v2, long standards used to monitor 5EU enrichment and recovery were added to total RNA (using a 1:10 ratio of 5EU-containing standard to non-5EU-containing standard), and samples were click labeled as above in reactions with 2.5 µg/µL RNA. For samples from the cell line 1 time course, click reactions were performed with 500, 500, 250, 200, or 100 µg total RNA for the 40 min, 1 h, 2 h, 4 h, or 8 h samples. For samples from the cell line 2 time course, click reactions were performed with 800, 525, 350, or 200 µg total RNA for the 40 min, 1 h, 2 h, or 4 h, respectively. For both cell lines, the steady-state samples did not undergo click reactions or pull-down.

### Purification of biotinylated RNA

For RNA-seq, Dynabeads MyOne Streptavidin C1 beads (ThermoFisher) for each set of samples were combined and batch washed, starting with 200 µL of beads per reaction. Beads were washed twice with 1X B&W buffer (5 mM Tris-HCl, pH 7.5, 0.5 mM EDTA, 1 M NaCl and 0.005% Tween-20), twice with solution A (0.1 M NaOH, 50 mM NaCl), twice with solution B (0.1 M NaCl), and then twice with water, using for each wash a volume equal to that of the initial bead suspension. Following the last wash, beads were resuspended in an initial bead volume of 1X high salt wash buffer (HSWB, 10 mM Tris-HCl, pH 7.4, 1 mM EDTA, 0.1 M NaCl, 0.01% Tween-20) supplemented with 0.5 µg/mL yeast RNA (ThermoFisher) and incubated at room temperature for 30 min with end-over-end rotation, again using a volume equal to that of the initial bead suspension. Beads were then washed three times with 200 µL 1X HSWB per reaction and split for each reaction during the last wash. After the wash was removed, sample RNA resuspended in 200 µL 1X HSWB was added to blocked beads and incubated with end-over-end rotation at room temperature for 30 min. Beads were washed twice with 800 µL 50°C water, incubating at 50°C for 2 min for each wash, and then twice with 800 µL 10X HSWB. RNA was eluted from beads by incubating with 200 µL 0.5 M tris(2-carboxyethyl)phosphine (TCEP, Sigma-Aldrich) at 50°C for 20 min with end-over-end rotation. The initial eluate was collected, and beads were resuspended in 150 µL water and eluted again, combining the two eluates for each sample. RNA from the eluate was then ethanol precipitated using linear acrylamide as a carrier.

Purifications of non-fragmented RNA were performed as above, except bead volumes were adjusted based on estimates of the amount of labeled RNA in each sample. For the cell line 1 samples, 292, 431, 410, 598, and 500 µL of beads were used for the 40 min, 1 h, 2 h, 4 h, and 8 h samples, respectively. For the cell line 2 samples, 467, 452, 575, and 598 µL streptavidin beads were used for the 40 min, 1 h, 2 h, and 4 h samples, respectively.

Pilot experiments designed to optimize the 5EU biotinylation and purification confirmed that RNAs containing at least one 5EU could be purified efficiently, with over 80% of a model RNA substrate containing a single 5EU becoming biotinylated in a 1 h reaction (Eisen et al., 2019). This high reaction efficiency was important for the RNA-seq samples, as RNA fragments from these libraries, generated to match ribosome-profiling samples (Eisen et al., 2019), were only ∼30 nt long and estimated to typically contain at most a single 5EU. Indeed, for each of the three protocols, which started with either full-length RNA (PAL-seq) or fragmented RNA (RNA-seq), metabolically labeled RNA was substantially enriched above background (Eisen et al., 2019).

### PAL-seq v2

This method starts with the same mRNA workup as PAL-seq v1 (Subtelny et al., 2014), except the design of the 3′ adapter allows for ligation to tails ending with a uridine nucleotide, as implemented in an improved version of TAIL-seq (Lim et al., 2016). PAL-seq v2 also includes a primer-extension reaction that occurs on the Illumina flowcell, with the goal of extending the sequencing primer all of the way through the poly(A) tail, so that the first sequencing read identifies both the mRNA and its cleavage-and-polyadenylation site, as in PAL-seq v1 (Subtelny et al., 2014). The poly(A)-tail length is then measured by direct sequencing of the poly(A) tail, as in TAIL-seq (Chang et al., 2014) (Figure S1A).

We used RNA standards of defined tail lengths to monitor library preparation, sequencing, and the computational pipeline for improved versions of PAL-seq and our implementation of TAIL-seq. Depletion of long-tailed sequences was the most prevalent source of measurement error. For TAIL-seq, this depletion seemed highly dependent on the sequencing protocol, with the best results obtained on a HiSeq machine in high-output mode using the v3 reagent kit.

Steady-state RNA (25 µg of unselected RNA from the 40 min sample) or half of the RNA eluted from each 5EU-selected sample was used to prepare PAL-seq libraries. Tail-length standard mixes (1 ng of set 1 and 2 ng of set 2 for each 5EU-selected sample, and twice these amounts for the steady-state sample), and trace 5′-radoiolabeled marker RNAs (Table S3) were added to each sample to assess tail-length measurements and ligation outcomes, respectively. Polyadenylated ends including those with a terminal uridine were ligated to a 3′-biotinylated adapter DNA oligonucleotide (1.8µM) in the presence of two splint DNA oligonucleotides (1.25µM and 0.25µM for the U and A-containing splint oligos, respectively, Table S3) using T4 Rnl2 (NEB) in an overnight reaction at 18°C. Following 3′-adapter ligation the RNA was extracted with phenol– chloroform (pH 8.0), precipitated, resuspended in 1X RNA T1 sequence buffer (ThemoFisher), heated to 50°C for 5 min and then put on ice. RNase T1 was then added to a final concentration of 0.006 U/µL, and the reaction was incubated at room temperature for 30 min, followed by phenol–chloroform extraction and RNA precipitation. Precipitated RNA was captured on streptavidin beads, 5′ phosphorylated, and ligated to a 5′ adapter as described (Subtelny et al., 2014) but using a modified 5′ adapter sequence (Table S3). Following reverse transcription using SuperScript III (Invitrogen) with a barcode-containing DNA primer, cDNA was purified as described (Subtelny et al., 2014), except a 160–810 nt size range was selected. Libraries were amplified by PCR for 8 cycles using Titanium Taq polymerase according to the manufacturer’s protocol with a 1.5 min combined annealing/extension step at 57°C. PCR-amplified libraries were purified using AMPure beads (Agencourt, 40 µL beads per 50 µL PCR, two rounds of purification) according to the manufacturer’s instructions.

The use of a splinted ligation of the 3′ adapter to the poly(A) tail had the advantage of specifically ligating to mRNAs without the need to deplete ribosomes or other abundant RNAs. However, this approach was not suitable for acquiring measurements for mRNAs with tails that were either very short (< 8 nt) or extended by more than one uridine, because such tails would ligate less efficiently (or not at all) when using a splinted ligation to the 3′ adapter. To account for these mRNAs with either very short or highly modified tails, we implemented a protocol that used single-stranded (ss) ligation and different mRNA enrichment steps to prepared libraries from steady-state RNA isolated from each of the two cell lines. For each sample, 5 µg of total RNA was depleted of rRNA using RiboZero Gold HMR (Illumina) and further depleted of the 5.8s rRNA by subtractive hybridization. Subtractive hybridization was performed by mixing 2x SSC buffer (3M sodium chloride, 300mM sodium citrate, pH 7.0), total RNA, and 4.8µM of each 5.8s subtractive-hybridization oligo (Table S3) in a 50 µL reaction, heating the reaction to 70°C for 5 min, then cooling it at 1°C/min to 37°C to anneal the oligos to the RNA. During this cooling, 250 µL of Dynabeads MyOne Streptavidin C1 beads per sample (ThermoFisher) were washed twice with 1X B&W buffer (5 mM Tris-HCl, pH 7.5, 0.5 mM EDTA, 1 M NaCl and 0.005% Tween-20), twice with solution A (0.1 M NaOH, 50 mM NaCl), twice with solution B (0.1 M NaCl), and then resuspended in 50 µL of 2X B&W buffer. After cooling, the entire 50µL RNA/oligo mixture was added to 50 µL of washed beads, then incubated at room temperature for 15 min with end-over-end rotation. The sample was then magnetized and the supernatant was withdrawn and precipitated by adding 284 µL of water, 4 µL of 5 mg/mL linear acrylamide, and 1 mL of ice-cold 96% ethanol. After resuspension, RNA was ligated to a 3′ adapter containing four random-sequence nucleotides and an adenylyl group at its 5′ end (Table S3) in a 70 µl reaction containing 10 µM adapter, 1X T4 RNA Ligase Reaction Buffer (NEB), 20 U/µL T4 RNA Ligase 2 truncated KQ (NEB), 0.3 U/µL SUPERaseIn (ThermoFisher), and 20% PEG 8000. The reaction was incubated at 22°C overnight and then stopped by addition of EDTA (3.5 mM final after bringing the reaction to 400 µL with water). RNA was phenol–chloroform extracted, precipitated, and subsequent library preparation was as for the splinted-ligation libraries.

PAL-seq v2 libraries were sequenced on an Illumina HiSeq 2500 operating in rapid mode. Hybridization mixes were prepared with 0.375 fmol PCR-amplified library that had been denatured with standard NaOH treatment and brought to a final volume of 125 µL with HT1 hybridization buffer (Illumina, 3 pM library in final mix). Following standard cluster generation and sequencing-primer hybridization, two dark cycles were performed for the splint-ligation libraries (i.e., two rounds of standard sequencing-by-synthesis in which imaging was skipped), which extended the sequencing primer by 2 nt, thereby enabling measurement of poly(A) tails terminating in non-adenosine bases. For the direct-ligation libraries, six dark cycles were performed instead of two, which extended the sequencing primer past the four random-sequence nucleotides in the 3′ adapter and then the last two residues of the tail.

Following the two dark cycles, a custom primer-extension reaction was performed on the sequencer using 50 µM dTTP as the only nucleoside triphosphate in the reaction. To perform this extension, the flow cell temperature was first set to 20°C. Then, 120 µL of universal sequencing buffer (USB, Illumina) was flowed over each lane, followed by 150 µL of Klenow buffer (NEB buffer 2 supplemented with 0.02% Tween-20). Reaction mix (Klenow buffer, 50 µM dTTP, and 0.1 U/µL Large Klenow Fragment, NEB) was then flowed on in two aliquots (150 µL and 100 µL). The flow-cell temperature was then increased to 37°C at a rate of 8.5°C per min and the incubation continued another 2 min after reaching 37°C. 150 µL of fresh reaction mix was then flowed in, and following a 2 min incubation, 75 µL of reaction mix was flowed in eight times, with each flow followed by a 2 min incubation. The reaction was stopped by decreasing the flow cell temperature to 20°C, flowing in 150 µL of quench buffer (Illumina HT2 buffer supplemented with 10 mM EDTA) and then washing with 75 µL of HT2 buffer. The flow cell was prepared for subsequent sequencing with a 150 µL and a 75 µL flow of HT1 buffer (Illumina). 50 cycles of standard sequencing-by-synthesis were then performed to yield the first sequencing read (read 1). XML files used for this protocol are provided at https://github.com/kslin/PAL-seq.

The flow cell was stripped, a barcode sequencing primer was annealed, and seven cycles of standard sequencing-by-synthesis were performed to read the barcode. The flow cell was then stripped again, and the same primer as used for read 1 was hybridized and used to prime 250 cycles of standard sequencing-by-synthesis to generate read 2. Thus, each PAL-seq tag consisted of three reads: read 1, read 2, and the indexing (barcode) read. For cases in which a tag corresponded to a polyadenylated mRNA, read 1 was the reverse complement of the 3′ end of the mRNA immediately 5′ of the poly(A) tail and was used to identify the mRNA and cleavage-and-polyadenylation site of long-tailed mRNAs. The indexing read was used to identify the sample, and read 2 was used to measure poly(A)-tail length and identify the mRNA and cleavage-and-polyadenylation site of short-tailed mRNAs. The intensity files of reads 1 and 2 were used for poly(A)-tail length determination, along with the Illumina fastq files.

### PAL-seq v2 data analysis

Tail lengths for the splinted-ligation data were determined using a Gaussian hidden Markov model (GHMM) from the python2.7 package ghmm (http://ghmm.org/), analogous to the model used in TAIL-seq (Chang et al., 2014) and described in the next paragraph. Read 1 was mapped using STAR (v2.5.4b) run with the parameters ‘-- alignIntronMax 1 --outFilterMultimapNmax 1 --outFilterMismatchNoverLmax 0.04 -- outFilterIntronMotifs RemoveNoncanonicalUnannotated --outSJfilterReads’, aligning to an index of the mouse genome built using mm10 transcript annotations that had been compressed to unique instances of each gene selecting the longest transcript and removing all overlapping transcripts on the same strand (Eichhorn et al., 2014). The genome index also included sequences of the quantification spikes and the common portion of the poly(A)-tail length standards. The sequences that identified each RNA standard (the last 20 nt of each standard sequence, Table S3) were not aligned using STAR. Instead, the unix program grep (v2.16) was used to determine which reads matched each standard (allowing no mismatches), and these reads were added to the aligned reads from the STAR output. Tags corresponding to annotated 3′ UTRs of mRNAs were identified using bedtools (v2.26.0), and if the poly(A)-tail read (read 2) contained a stretch of ≥ 10 T residues (the reverse complement of the tail) in an 11-nt window within the first 30 nt, this read was carried forward for GHMM analysis. If read 2 failed to satisfy this criterion but began with ≥ 4 T residues, the tail length was called based on the number of contiguous T residues at the start of read 2; by definition, these tails were < 10 nt and thus easily determined by direct sequencing.

For each read 2 that was to be input into the GHMM a ‘T signal’ was first calculated by normalizing the intensity of each channel for each cycle to the average intensity of that channel when reading that base in read 1 and then dividing the thymidine channel by the sum of the other three channels. Sometimes a position in a read would have a value of 0 for all four channels. A read was discarded if it contained more than five such positions. Otherwise, the values for these positions were imputed using the mean of the five non-zero signal values upstream and downstream (ten positions total) of the zero-valued position. A three-state GHMM was then used to decode the sequence of states that occurred in read 2. It consisted of an initiation state (state 1), a poly(A)-tail state (state 2), and a non-poly(A)-tail state (state 3). All reads start in state 1. From state 1 the model can remain in state 1 or transition to state 2. From state 2 the model can either remain in state 2 or transition to state 3. The model was initialized with the following transition probabilities:

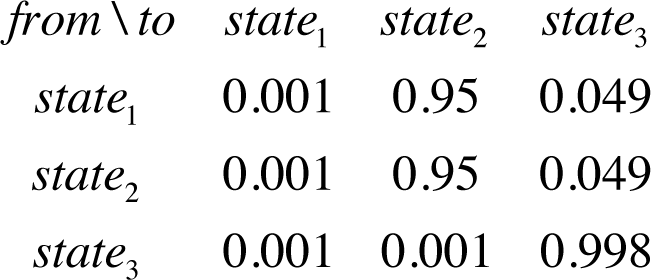

The initial emissions were Gaussian distributions with means of 100, 1, and –1 and variances of 1, 0.25 and 0.25, respectively. In general, the emission Gaussians for the model corresponded to the logarithm of the calculated T signal at each sequenced base in read 2. The initial state probabilities were 0.998, 0.001, and 0.001 for states 1, 2 and 3, respectively.

After initializing the model, unsupervised training was performed on 10,000 randomly selected PAL-seq tags, and then the trained model was used to decode all tags, with the number of state 2 cycles reporting the poly(A)-tail length for a tag. Only genes with ≥ 50 poly(A)-tail length measurements were considered for analyses involving mean poly(A)-tail lengths.

### Analysis of PAL-seq data from the ss-ligation protocol

To account for mRNAs with very short tails or extensive terminal modifications, we implemented a version of PAL-seq that did not use splinted ligation. Tail lengths from these ss-ligation datasets, acquired for steady-state samples from both cell lines, were determined using a modified version of the PAL-seq analysis pipeline written for python3. The T-signal in this pipeline was modified to allow more accurate quantification of 0-length tails. Instead of normalizing the intensity of each channel for each cycle to the average intensity of that channel when reading that base in read 1, the intensity of each channel was normalized to the average intensity of the channels for the other three bases in read 1. The intensity of the T channel was then divided by the sum of the other channel intensities to calculate the T signal, and tails were called using the hmmlearn package (v0.2.0). Tags representing short tails, including short tails that ended with many non-A residues, were identified as those for which read 1 and read 2 mapped to the same mRNA 3′ UTR (usually ∼4% of the tags). Tail lengths for these tags were called without the use of the GHMM. Instead, their tail lengths were determined by string matching, allowing any number of untemplated U residues but no more than two G or C residues to precede the A stretch. Tags not identified as representing short-tails were analyzed using the GHMM, excluding from further analysis occasional outliers determined by the GHMM to have tails ≤ 8 nt.

Most of the tags that had either only a very short tail or no tail did not correspond to mRNA cleavage-and-polyadenylation sites. Therefore, to be carried forward in our analysis, short-tailed tags were required to have a 3′-most genome mapping position (as determined from read 1 but requiring that read 2 also map uniquely to the same 3′ UTR) that fell within a 10 nt window of a PAL-seq–annotated cleavage-and-polyadenylation site.

Although the single-stranded ligation protocol provided the opportunity to account for mRNAs with very short or highly modified tails, examination of the recovery of internal standards indicated that tags representing longer tails (≥ 100 nt) were not as well recovered in the datasets in which we implemented ss ligation. Therefore, for steady-state samples from each cell line, we generated composite tail-length distributions in which the ss-ligation dataset contributed to the distribution of tails < 50 nt, and the splint-ligation dataset contributed to the distribution of tails ≥ 50 nt. For example, *Slc38a2* had 635 standard PAL-seq tags, 169 of which (∼27%) had tails < 50 nt, and this same gene had 703 ss-ligation PAL-seq tags, 393 of which (∼56%) had tails < 50 nt. The composite tail-length distribution replaced the 169 short-tailed splint-ligation PAL-seq tags with the 393 short-tailed ss-ligation PAL-seq tags, normalizing the latter cohort by a scaling factor. This scaling factor was determined from the ratio of the counts of the splint-ligation tags with tail lengths between 30–70 nt (135 tags) to the counts of the corresponding tags in the ss-ligation dataset (153 tags).

3′-end annotations were generated from PAL-seq tags with tails ≥ 11 nt, using an algorithm previously developed for data from poly(A)-position profiling by sequencing (3P-seq) (Jan et al., 2011). Each PAL-seq read 1 that mapped (with at least 1 nt of overlap) to an annotated 3′ UTR (Eichhorn et al., 2014) was compiled by the genomic coordinate of its 3′-most nucleotide. The position with the most mapped reads was annotated as a 3′ end. All reads within 10 nt of this end (a 21 nt window) were assigned to this end and removed from subsequent consideration. This process was repeated until there were no remaining 3′ UTR-mapped reads. For each gene, the 3′-end annotations were used in subsequent analyses if they accounted for ≥ 10% of the 3′ UTR-mapping reads for that gene.

Documentation and code to calculate and analyze T signals and determine tail lengths are available for both the splint-ligation and ss-ligation pipelines at https://github.com/kslin/PAL-seq.

### TAIL-seq library preparation, sequencing, and analysis

The 2 h time-interval TAIL-seq used for comparison with PAL-seq was prepared using the same library cDNA as was used for PAL-seq v2 libraries, but amplifying the library using different primers (Table S3). Amplification and purification were as for PAL-seq v2. Samples were sequenced with either a paired-end 50-by-250 run (2 h time-interval sample) using a HiSeq 2500 operating in normal mode using a v3 kit. Other Illumina sequencing chemistries (including v1, v2, and v4 kits run in rapid and normal modes) did not yield accurate tail-length measurements when used in paired-end mode. Analysis was as described for PAL-seq v2, except a five-state GHMM was used (Chang et al., 2014) to accommodate the difference in the nature of the T-signal output imparted by the different mode of sequencing. The five states were an initiation state, a poly(A) state, a poly(A) transition state, a non-poly(A) transition state, and a non-poly(A) state.

### RNA-seq

Fragmented poly(A)-selected RNAs were supplemented with three short quantification standards (Table S3), and then ligated to adapters, reverse-transcribed, and amplified to prepare the RNA-seq and ribosome-profiling libraries, respectively (Subtelny et al., 2014). These libraries were sequenced on an Illumina HiSeq 2500. For all RNA-seq data, only reads mapping to ORFs of annotated gene models (Eichhorn et al., 2014) were considered, excluding the first 50 nt of each ORF, which was implemented to match ribosome-profiling data of a contemporary study examining the effects of miRNAs (Eisen et al., 2019). A cutoff of ≥ 10 reads per million mapped reads (RPM) was applied to each sample.

### Calculation of mRNA half-lives

Half-lives were estimated independently from both RNA-seq data and PAL-seq tag abundance. Prior to half-life fitting, mRNA abundances were normalized across time intervals based on the quantification standards added to each sample prior to library preparation.

Half-lives were determined by fitting to the equation

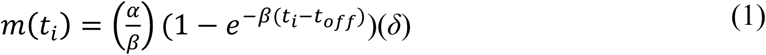

in the case of the continuous-labeling experiment, or to the equation

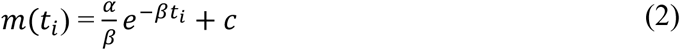

in the case of the transcriptional shutoff experiment, where *m*(*t_i_*) is the expression of an mRNA at a given time *i*, *α* is the rate of mRNA production, *β* is the rate of mRNA degradation, *t_off_* is a global time offset, *δ* is a global scaling parameter to adjust the steady-state time point, and *c* is a baseline for the final expression of the gene in the transcriptional-shutoff experiment. Because the quantification standards were not applicable to the steady-state sample, the steady-state sample was normalized by a globally-fitted constant (setting *t_i_* to 100 h for this time interval).

Because the half-life fitting for the continuous-labeling experiment required the global parameters *t_off_* and *δ*, half-lives for all genes needed to be fit simultaneously. Accordingly, we minimized the least-square errors loss function (*L*_2_).

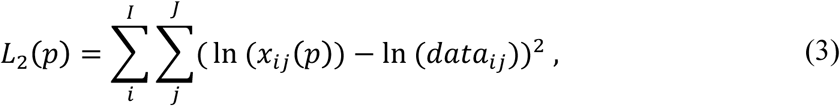

for the simulated number of normalized tags *x* at time point *i* for gene *j*. The total number of time points and genes are denoted by *I* and *J*, respectively. *L*_2_ depends on the parameters *p* (*α_i_*_,*j*_, *β_i_*_,*j*_, *t_off_*, and *δ*). The optimization for *α_i_*_,*j*_, *β_i_*_,*j*_, *t_off_*, and *δ* was performed using the L-BFGS-B method in the optim function in R.

To increase the efficiency of the optimization, we also implemented an analytical gradient for this model. This gradient computed the quantity 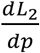 which, when passed to the optimizer, decreased the number of iterations required to minimize the loss. This quantity was computed for each of the parameters as follows

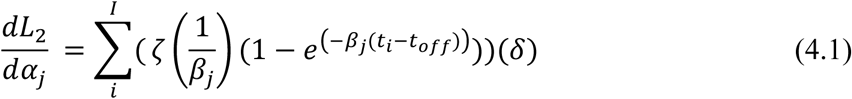

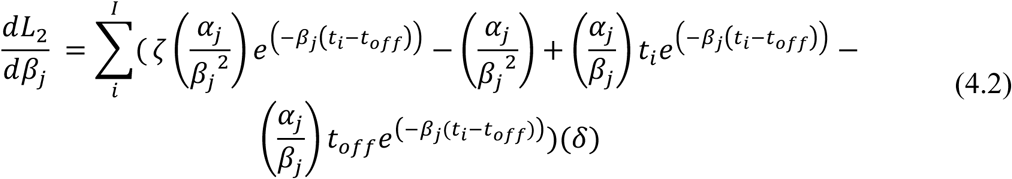

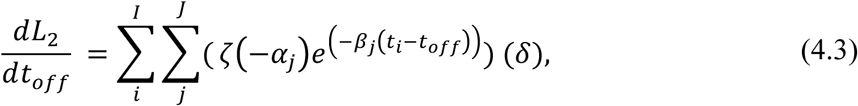

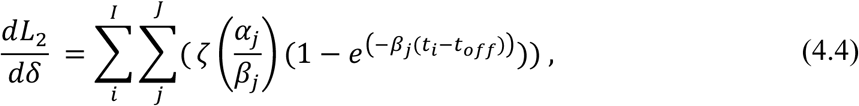

where *ζ* is the first component of the derivative of the loss function

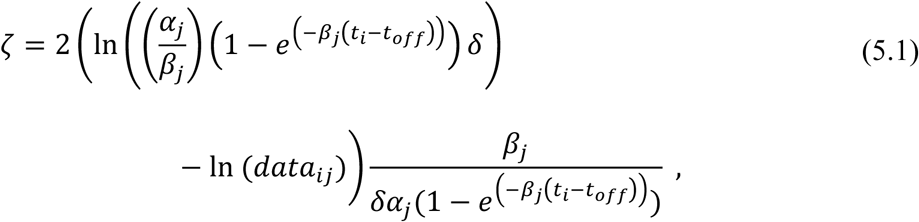

and *δ* is further defined by the piecewise function

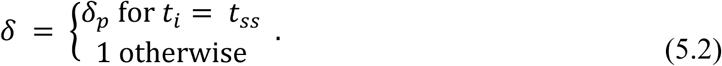

Ranges of rates constants fit to this exponential model and the subsequent deadenylation model were truncated to reflect the lack of confidence in values of and differences between extreme outliers. Half-live values were truncated to fall between 6 min and 100 h, deadenylation rate constants were truncated to fall between 0.03 and 30 nt/min, decapping rate constants at 20 nt were truncated to fall between 0.003 and 3 min^−1^, and production rate constants were truncated to fall between 10^−8^ and 10^−5^ min^−1^. All calculations (including correlations) were performed on the non-truncated values.

### Model of mRNA production, deadenylation, and decay

The model of mRNA production, deadenylation, and decapping (decay) was a system of differential equations

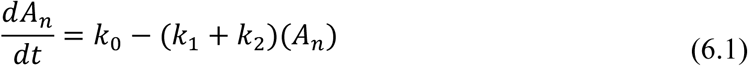

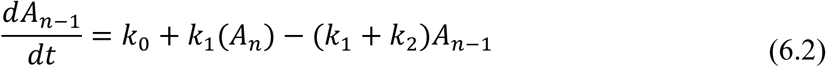

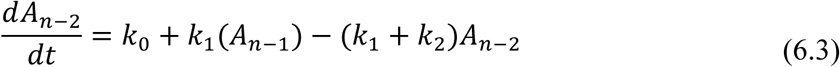

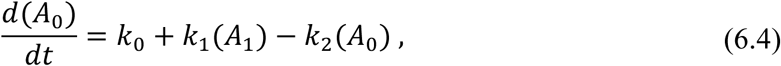

where *A_n_* is an mRNA with tail length *n*, and *k*_0_, *k*_1_ and *k*_2_ are rate constants that describe the production, deadenylation and decapping rates, respectively. The final deadenylation product (*A*_0_) has a deadenylation rate constant of zero, as it has no tail. The rate constants *k*_0_ and *k*_2_ are themselves functions of tail length (*l*), specified by the respective negative binomial and logistic functions

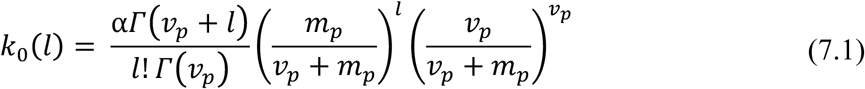

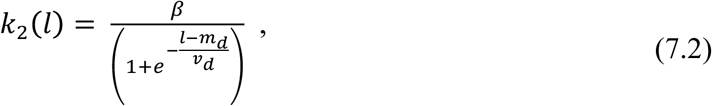

where the parameters *α*, *β*, *v_p_*, *m_p_*, *m_d_*, *v_d_*, are fitted parameters. The parameters *α* and *β* are scaling terms for production and decapping distributions, respectively. The parameters *m_p_* and *m_d_* describe the expected value of those distributions, and *v_p_* and *v_d_* describe the spread.

Equations (6) were re-written as a linear, time-invariant (LTI) system (Dahleh et al., 2004)

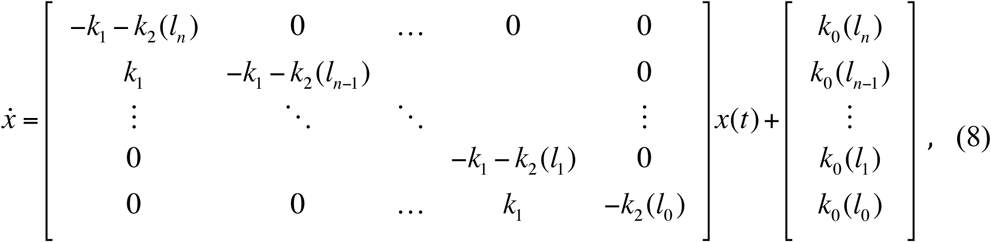

or, more succinctly, as

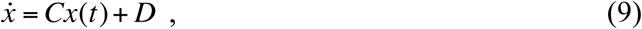

where the coefficient matrix, *C*, is specified by the coefficients of the differential equations (6), and the source vector *D* is specified by the production rate. *C* is a bidiagonal 251×251 matrix, whereas *D*, *x(t)*, and *ẋ* are 251×1 vectors. In the case of the continuous-labeling experiment, *x t* = 0 = **0**. The transcriptional-shutoff experiment begins with *x t* = – 1 h = **0**, but *x t* = 0 is determined by the values of the system after 1 h of simulation.

Equation (9) has the analytical solution

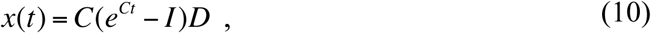

where *I* is the identity matrix and *e^Ct^* is the matrix exponential of the coefficient matrix scaled by time. Both this analytical solution and numerical integrators (which do not require an analytical solution) can be used to compute the result. We found that numerical stability and computational efficiency were optimal when using the LSODE solver with parameters set for a banded Jacobian matrix in the deSolve package (v1.21) of R, with the model written in C and dynamically loaded into R. Increasing the number of allowed tail-length states from 250 to 300 had little effect on the resulting fitted rate constants but greatly increased computation time.

The model yielded abundances for each tail-length isoform at each time interval, using seven parameters for each gene, three of which were shared across all of the genes (Table S2). From these abundances, the residual sum of squares was computed from the corresponding standard-normalized PAL-seq datasets. Although tail lengths ≥ 250 nt were modeled, measurements for these lengths were not available from PAL-seq v2, and thus were excluded from the fitting. Likewise, tail-lengths < 20 were modeled for all time intervals, but because the abundance of tail lengths < 20 nt was only available for steady state, these lengths were excluded from fitting all but the steady-state interval. As a result, for the continuous-labeling experiment using cell line 1, parameters for each gene were fit to 1400 data points (230 tail lengths × 5 time intervals + 250 for the steady-state), and for the experiment using the cell line 2, parameters for each gene were fit to 1170 data points. The optimization was performed using the L-BFGS-B method in the optim function of R, or, in the case of the global fitting, using the L-BFGS-B method in the NLopt package (v1.0.4) of R.

A simple *L*_2_ loss function skewed the fits to the time intervals that had larger values. A common solution to this problem is to fit to log-transformed values, but because our data were sparse, with many tail-length positions having zero tags, pseudo-counting to allow log-space fitting resulted in poor fits. Therefore, residuals were variance weighted using the loss function

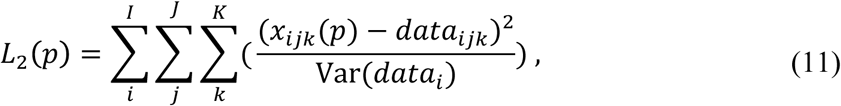

where *i*, *j*, and *k* are the time-interval, gene, and tail length with *I*, *J* and *K* as the maximal values of time-intervals, genes, and tail lengths. The variance of a dataset at a single time-interval is given by Var(*data_i_*).

The model is constrained by the 0-tail-length species, which builds up when decapping is slow with respect to deadenylation. Such a buildup was observed in the steady-state tail-length distribution of short-lived mRNAs but occurred primarily between 0 and 20 nucleotides (Figure 6C). Because of this discrepancy, a composite residual was calculated for the model and the data. Abundances for tails < 20 nt were averaged and this average was used to replace the abundances for each tail length < 20 for the steady-state data. In addition, when comparing the associated short tails from the data and the model, the residuals for tails < 20 were weighted by either 6- or 5-fold (cell lines 1 and 2, respectively) to account for opting not to fit to measurements for tails < 20 nt in the non-steady–state samples.

As with gene-specific parameters, global parameters *v_p_*, *m_d_*, and *v_d_* were fit using pre-steady-state measurements of tails ranging from 20–249 nt. The composite steady-state tail-length distributions of Figure 2A were also used, which constrained buildup of short-tailed mRNAs. Fitting was performed on subsets of 100 genes (selected randomly without replacement from genes with composite steady-state distributions, yielding 22 and 15 subsets for cell lines 1 and 2, respectively), including *v_p_*, *m_d_*, and *v_d_* in the parameter vector. Median values of the global parameters (Table S1) were then used to fit each gene-specific parameter.

### Bootstrap analysis

Tags in the cell line 1 PAL-seq dataset were resampled 10 times with replacement and assigned to a gene and tail length based on a multinomial probability distribution generated from the counts for each tail length in the original dataset. These resampled datasets were then used for background subtraction, global parameter determination, and model fitting.

### Background subtraction and normalization for PAL-seq data

Although the efficacy of the 5EU purification enabled efficient enrichment of labeled RNAs at short time intervals (Eisen et al., 2019), we also modeled and corrected for residual background caused by non-specific binding of the unlabeled RNA to the streptavidin beads (Figure S2F).

We designed our background model under the assumption that the background in the time courses stems primarily from the capture of a fixed amount of non-5EU labeled mRNA during the 5EU purification. Accordingly, we subtracted a fraction (0.3%) of the steady-state data from each continuous-labeling dataset. This fraction of input sample was chosen such that at 40 min long-lived genes (half-life ≥ 8 h) had no mRNAs with tail lengths ≤ ∼100 nt on average, but short-lived genes (half-life ≤ 30 min) were unaffected (Figure S2F). Likewise, we subtracted standard-normalized time-interval–matched input data from each transcriptional-inhibition dataset, as actD influenced which unlabeled cellular mRNAs were available to contribute to the background. The fraction of each input sample to subtract was chosen such that at 0 h long-lived genes (half-life ≥ 8 h) had no mRNAs with tail lengths ≤ ∼100 nt on average, but short-lived genes (half-life ≤ 30 min) were unaffected. Genes were included in the final background-subtracted set only if the sum of their background-subtracted tag counts was ≥ 50 tags.

After background subtraction, PAL-seq datasets were scaled to each time interval by matching the total number of background-subtracted tags for all genes at all tail lengths to the total number of tags for all genes for the corresponding time interval in the RNA-seq data. The scaled PAL-seq data were then used to compute half-lives for each gene, scaling the steady-state sample using a globally fitted constant.

### ActD treatment

Cell line 2 was cultured as in the continuous-labeling experiments. We prepared 2, 2, 2, 3, and 4 500 cm^2^ plates for the 0, 1, 3, 7 and 15 h time intervals, respectively. 5EU (400 µM final) was added to each plate (with one non-5EU plate prepared in parallel), and after 1 h actD (5 µg/mL final concentration, Sigma-Aldrich) was added. Cells were harvested as described for the continuous-labeling experiments, except that a quantitative spike RNA containing 5EU and corresponding to the chloramphenicol-resistance gene sequence (Table S3) was added to the lysis buffer at a concentration of 0.57 ng/mL, or 2 ng/plate. This RNA was prepared using an in vitro transcription reaction as above, with a 5EUTP-to-UTP ratio of 1:20.

### Accession numbers

Raw and processed RNA-seq, PAL-seq, and TAIL-seq data is available at the GEO, accession number GSE134660. Code for configuring an Illumina HiSeq 2500 machine for PAL-seq and for calculation of tail lengths from PAL-seq or TAIL-seq data is available at https://github.com/kslin/PAL-seq.

## Supplemental Figure Legends

**Figure S1.**
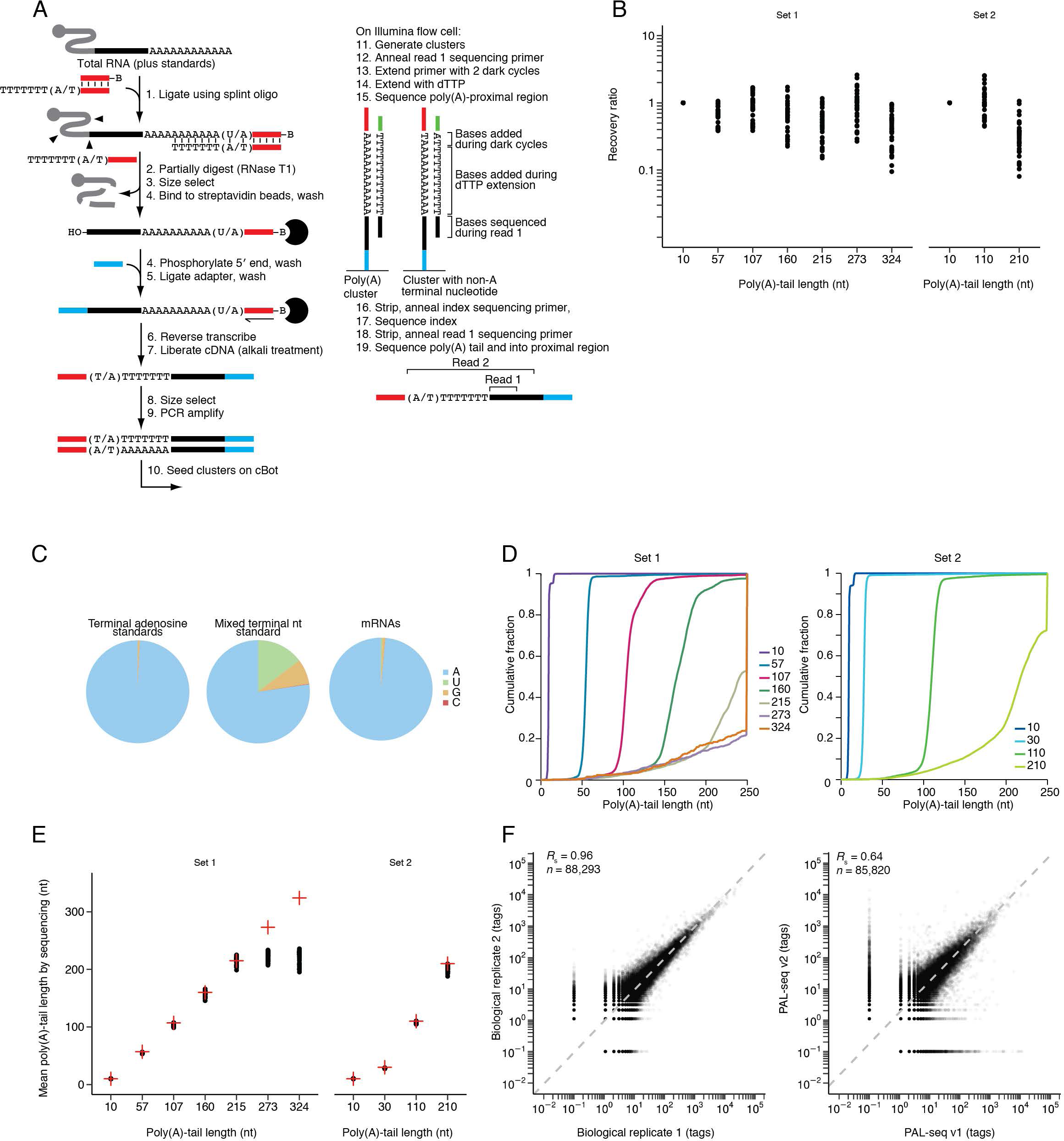
PAL-seq v2 Methodology and Benchmarking, Related to Figure 1. (A) Schematic of PAL-seq v2. The original version of PAL-seq (Subtelny et al., 2014) was modified to include an additional splint oligonucleotide capable of ligating to tails with a terminal U (step 1); two dark cycles prior to the primer-extension reaction (step 13), which prevented non-adenosine terminal residues from terminating the subsequent primer extension; primer extension through the tail with dTTP as the only nucleoside triphosphate (step 14); sequencing on a HiSeq machine, with the opportunity for multiplexing (steps 16 and 17); an additional read using the read 1 sequencing primer (read 2), which collected sequence and intensity information used to call poly(A)-tail lengths, as in TAIL-seq (Chang et al., 2014) (steps 18 and 19). (B) Recovery of RNA standards. Before preparing libraries, two sets of RNA standards were added to each of the 34 RNA samples analyzed by PAL-seq v2 in this study and an accompanying study (Eisen et al., 2019). Set 1 contained seven RNAs with different tail lengths, and set 2 contained four RNAs with different tail lengths (Table S3). For each set of standards, the relative abundance of each standard in the final sequencing output was compared its relative abundance in the initial standard mixture, and this recovery ratio is plotted for each sample on a log scale. The relative recovery of standards varied somewhat, with no systematic bias that would indicate substantial depletion of poly(A)-tails of certain lengths. The 30 nt standard from set 2 was excluded from this analysis because it is an equal mixture of four different standards that end in a terminal A, C, G or U (Table S3), which was added to assess the ability to detect tails with a terminal U, as described in the next panel. (C) Terminal nucleotide compositions of RNAs with tail measurements ≥ 5 nt. Libraries were prepared using a 5:1 mixture of splint oligos that would hybridize perfectly to either the 3′ end of RNAs ending in eight adenosines or the 3′ end of RNAs ending in seven adenosines followed by a terminal uridine, respectively. Left: Terminal nucleotide composition of PAL-seq v2 tags from the RNA standards for which poly(A) tails were prepared using poly(A) polymerase and ATP. These standards were expected to terminate exclusively with adenosine. Middle: Terminal nucleotide composition of PAL-seq v2 tags from the synthetic 30 nt standard, which was prepared with a tail designed to have an equal mixture of terminal A, C, G, or U. Although the splint oligonucleotides perfectly matched the versions ending A and U, the terminal U was somewhat depleted compared to the terminal A, and a substantial fraction of terminal G was also captured, perhaps due to wobble-pairing between the T in the splint and the terminal G in the standard. Right: Terminal nucleotide composition of PAL-seq v2 tags from mRNAs. (D) The tail length distributions of the synthetic RNA standards, as measured by PAL-seq v2. Plotted is the cumulative distribution of poly(A)-tail lengths for each standard in the steady-state sample from cell line 1. The poly(A)-tail lengths measured by polyacrylamide gel electrophoresis (Subtelny et al., 2014) are indicated (key). (E) Mean poly(A)-tail lengths of the two sets of synthetic standards, as measured by PAL-seq v2. For each standard, the mean tail length in each of the 34 samples in this study and an accompanying study (Eisen et al., 2019), as measured using PAL-seq v2, is plotted (black points). Also plotted is the mean tail length of each standard, as determined using denaturing gels (red crosses). Tails that exceeded 250 nt were not expected to be measured accurately, because their length exceeded the length of the sequencing read used to measure the tail. (F) Comparison of the frequencies in which UTR 3′ ends were called by biological replicates of PAL-seq v2 (left) or the two different versions of PAL-seq (right). Each point is the number of tags that mapped to a genomic position, adding a pseudo-count of 0.1 tags. Dashed lines represent equivalency, after accounting for different sequencing depths.

**Figure S2.**
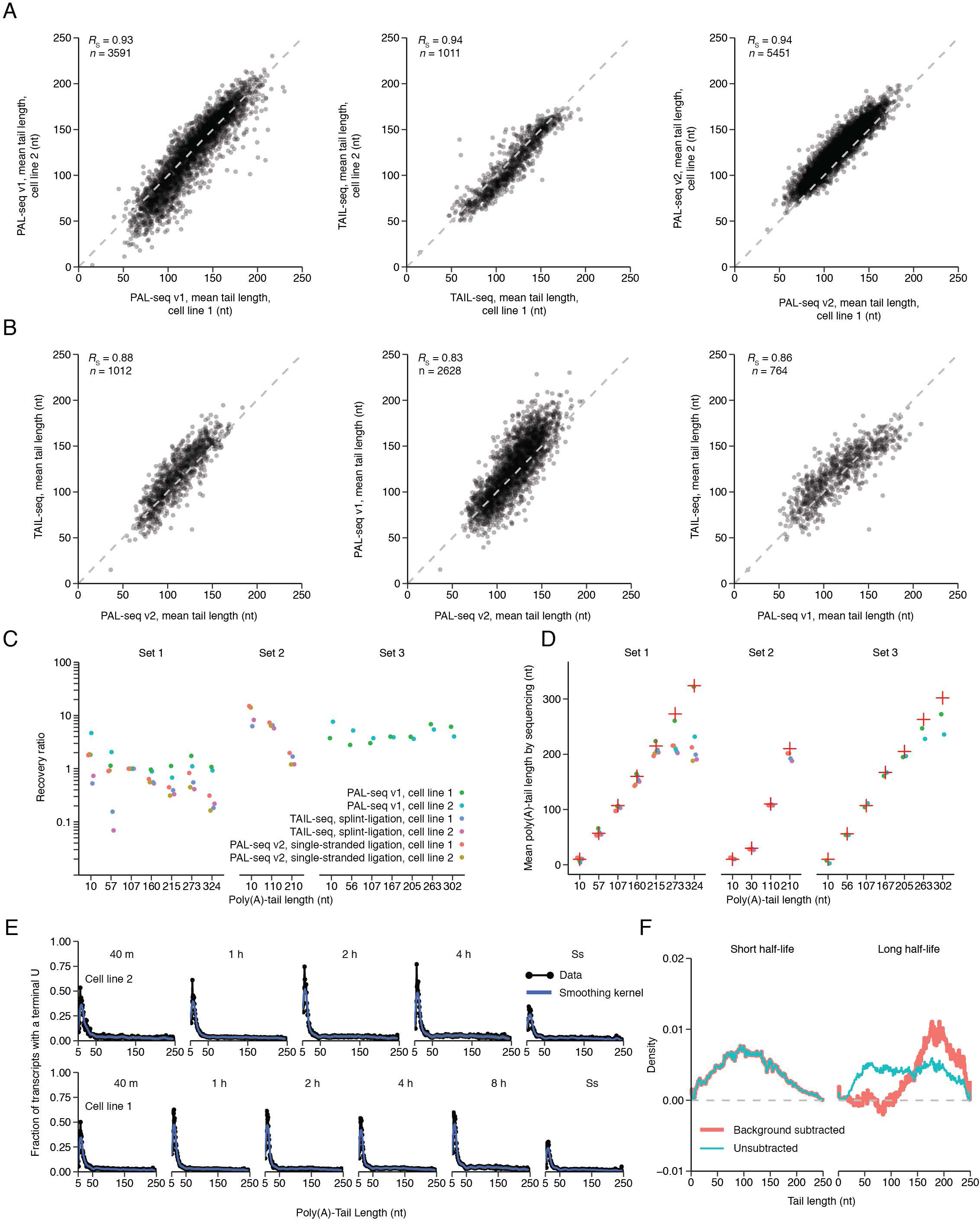
Reproducibility of PAL-seq v2, Related to Figure 1. (A) Comparisons of biological replicates for different library preparation and tail-profiling protocols. For each gene that passed a 50-tag cutoff, mean poly(A)-tail lengths after 2 h of continuous labeling are shown for PAL-seq (left panel), our implementation of TAIL-seq (Lim et al., 2016), (middle panel), or PAL-seq v2 (right panel). Whereas the RNA examined using TAIL-seq and PAL-seq v2 datasets was isolated using 5EU labeling, the RNA examined using PAL-seq v1 dataset was isolated using 4-thiouridine labeling. The 2 h time interval was chosen for this analysis because its broad range of average tail lengths made it most suitable for comparing the results of different methods (Figure 1C). The dashed line represents y = x. (B) Comparisons between different tail-profiling protocols. Compared are mean tail lengths generated by TAIL-seq and PAL-seq v2 (left panel), PAL-seq v1 and PAL-seq v2 (middle panel), and TAIL-seq and PAL-seq v1 (right panel). Otherwise, as in (A). (C) Recovery of tail standards in the PAL-seq v1, TAIL-seq (splinted ligation) and PAL-seq v2 steady-state (single-stranded ligation) datasets. For analyses of the recovery of tail standards in PAL-seq v2 (splint-ligation) datasets, see Figure S1B. All six libraries contained seven standards from standard set 1; the TAIL-seq libraries contained three standards from standard set 2; and the PAL-seq v1 library contained seven standards from standard set 3. Tail lengths of the standards as determined by polyacrylamide-gel electrophoresis (Subtelny et al., 2014) are indicated (key) and shown as x-axis labels. The 30 nt standard from set 2 was excluded from this analysis because it was an equal mixture of four different standards that ended in A, C, G or U (Table S3) and was added to assess the ability to detect tails with a terminal U. The relative abundances of the standards in the sequencing data were quantified and compared to their relative starting abundance, and this recovery ratio is plotted for all samples. The values of each library were normalized to the abundance of the 107 nt standard in set 1. (D) Mean tail lengths of the standards shown in (C) and the 30 nt standard. Otherwise, as in (C). For analysis of mean tail lengths of the standards in PAL-seq v2 (splint-ligation) datasets, see Figure S1E. (E) Uridylation frequency as a function of tail length. The fraction of single uridine residues at the 3′ end of mRNA-mapping tags is plotted as a function of tail lengths ≥ 5 nt (black) along with a LOESS smoothing kernel (blue, with 5^th^–95^th^ percent confidence intervals in grey) for either cell line 2 (top panels) or cell line 1 (bottom panels). These values were scaled by a factor of 5.23 to correct for the depletion of tags containing a terminal uridine, estimated from the ratio of tags mapping to the 30 nt standards terminating with either A or U, which had been added to the libraries at an equal molar ratio (Figure S1C). Uridine fractions corresponding to tail lengths ≥ 246 nt were combined into one bin at 246 nt. (F) Effects of background subtraction of PAL-seq data at the earliest (40 min) time interval. Distributions of the unsubtracted (blue) and background-subtracted (red) tail lengths for short-lived (half-life < 30 min, n = 293 for both unsubtracted and background subtracted) and long-lived (half-life > 8 h, n = 379) mRNAs. The background subtraction differentially affected the long half-life mRNAs, as these had a proportionally smaller amount of labeled relative to unlabeled RNA at short time intervals, and thus unlabeled RNAs contributed a larger fraction of their reads at these intervals.

**Figure S3.**
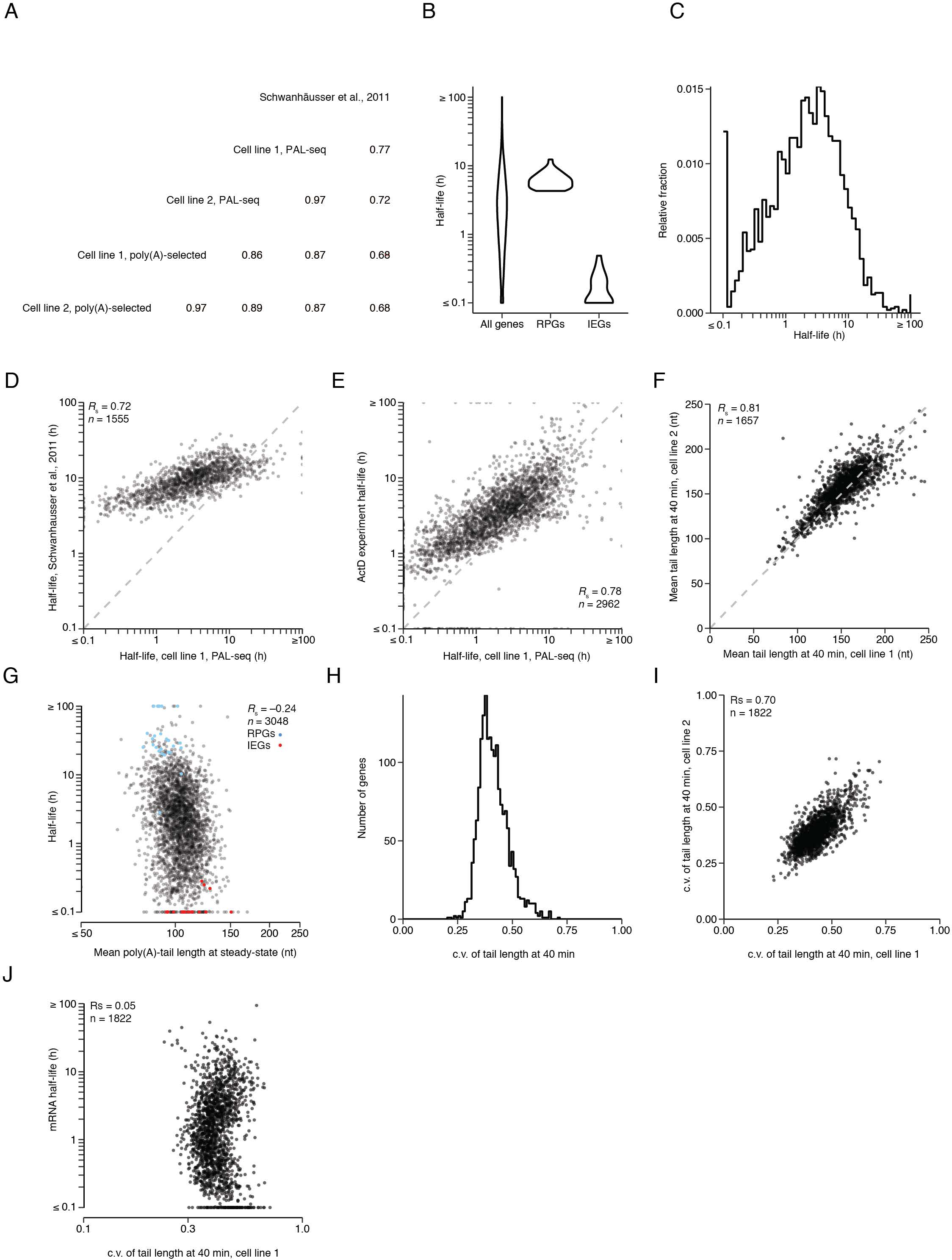
Half-life and Initial Tail-Length Measurements, Related to Figures 2 and 3. (A) Pairwise correlations (*R*_s_) of half-life measurements. n = 4485, 4748, 3048, 1743, and 4658 genes for cell line 1 poly(A)-selected, cell line 2 poly(A)-selected, cell line 1 PAL-seq, cell line 2 PAL-seq, and Schwanhäusser et al. 2011 samples, respectively. (B) Distributions of half-lives for mRNAs from all genes (n = 3048), ribosomal-protein genes (RPGs, n = 31), or immediate-early genes (IEGs, n = 19) (Tullai et al., 2007) obtained using PAL-seq data from cell line 1. (C) Distribution of mRNA half-lives obtained using PAL-seq data from cell line 1 (n = 3048). (D) Comparison of published half-life measurements (Schwanhäusser et al., 2011) with those obtained from 5EU continuous labeling. Dashed line is *y* = *x*. (E) Comparison of half-life measurements from the transcriptional-shutoff experiment and those obtained from the continuous-labeling experiment. Dashed line is *y* = *x*. (F) Comparison of mean poly(A)-tail lengths of mRNAs isolated from cell line 2 after 40 min of labeling with those isolated from cell line 1 after 40 min of labeling. Dashed line is *y* = *x*. (G) Relationship between half-life and mean steady-state tail length of mRNAs in 3T3 cells. Tail lengths and half-lives were determined using only standard PAL-seq data, in which the adapter oligo was appended to the tail using splinted ligation. Otherwise, as in Figure 2A. (H) The distribution of c.v. values of tail lengths after 40 min of labeling for mRNAs from each gene. Each c.v. value is the average of two biological replicates. (I) Comparison of c.v. values of tail lengths after 40 min of labeling between two biological replicates. (J) Relationship between mRNA half-life and c.v. values of tail lengths after 40 min of labeling. Each c.v. value is the average of two biological replicates.

**Figure S4.**
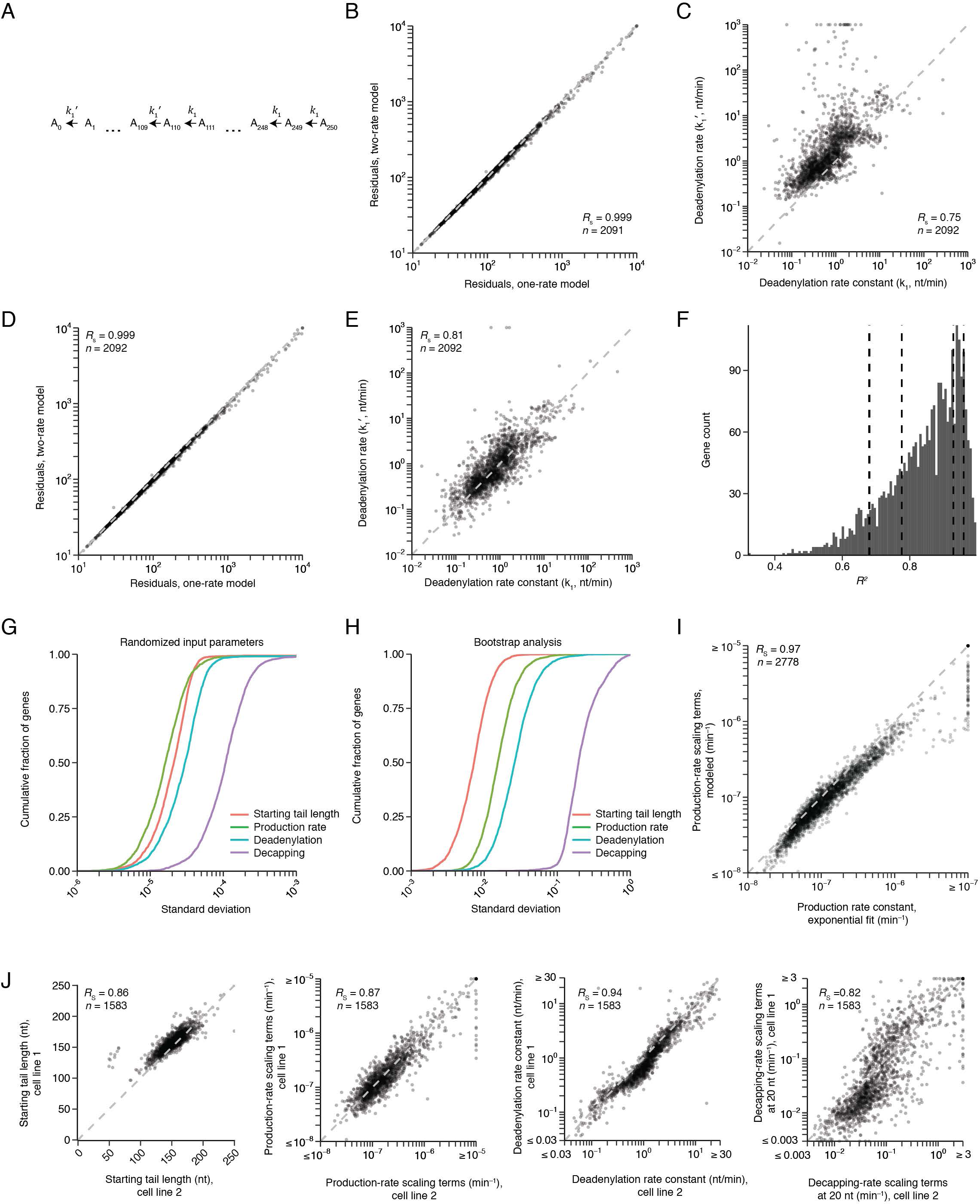
Model Development and Testing, Related to Figures 4 and 5. (A) Schematic of the model with two deadenylation rates. Deadenylation is parameterized with two rate constants, one that describes deadenylation of tail lengths > 110 nt (*k*_1_) and one that describes deadenylation of tails ≤ 110 nt (*k*_1_′). The transition between these rates is determined by a generalized logistic function with a transition parameter arbitrarily set to 1 (a sharp transition). Otherwise, as in Figure 4A. (B) Comparison of the residual sum of squares (RSS) between the model with two deadenylation rates (A) and the model with one deadenylation rate. Dashed line is *y* = *x*. (C) Relationship between the second (*k*_1_′) and the first (*k*_1_) deadenylation rate constant fit for mRNAs of each gene using the model in (A). Dashed line is *y* = *x*. (D) Comparison of a model with two deadenylation rates in which the transition between the rates occurs at a tail length of 150 nt and the model with one deadenylation rate. Otherwise, as in (B). (E) Relationship between the second (*k*_1_′) and the first (*k*_1_) deadenylation rate fit for mRNAs of each gene using the model in (D). Dashed line is *y* = *x*. (F) Distribution of *R^2^* values for all genes fit by the model (n = 2778). Dashed lines indicate the *R*^2^ values of the four genes shown in Figure 4B. (G) Analysis of the robustness of fitted rate constants to input parameter identities. The distributions of s.d. values of rate constants for all fitted genes over 10 rounds of fitting with varying input parameters are displayed as empirical cumulative distributions. The input parameters were randomly selected from a uniform distribution bounded by the 10^th^ to 90^th^ percentiles of rate constants of all genes during a previous round of fitting. Using an unbounded randomized parameter selection resulted in larger variation but also larger final residuals. The s.d. values for 90% of genes were less than 3.7 × 10^−5^, 3.5 × 10^−5^, 5.8 × 10^−5^, and 2.4 × 10^−4^ for rate constants for starting tail length, production, deadenylation, and decapping, respectively (with all parameters shown as s.d. of the log_10_ of the value). (H) Bootstrapping analysis of fitted rate constants. For each dataset, the total number of tags was resampled ten times based on a multinomial probability distribution specified by the original tag counts for every tail length position for each gene. These resampled datasets were then background subtracted and fit to the computational model. Shown for each fitted gene-specific parameter is the cumulative distributions of its c.v. values for all fitted genes. The s.d. for 90% of genes were less than 0.01, 0.03, 0.06, and 0.49, for rate constants for starting tail length, production, deadenylation, and decapping, respectively (with all parameters shown as s.d. of the log_10_ of the value). The greater variation observed for the decapping parameter reflected the relatively few data points effectively used for its fitting, as this parameter related primarily to the short-tailed region of the distribution. (I) Relationship between production rate from an exponential fit to the data and the production rate as determined by the computational model. Dashed line is *y* = *x*. (J) Model reproducibility across biological replicates. Plots show the relationship between mean starting tail lengths (*m_p_*, left panel), production-rate scaling terms (*α*, middle left panel), deadenylation rate constants (*δ*, middle right panel) or decapping-rate scaling terms (*β*, right panel) for mRNAs from the same genes in the two cell lines. Dashed line is *y* = *x*.

**Figure S5.**
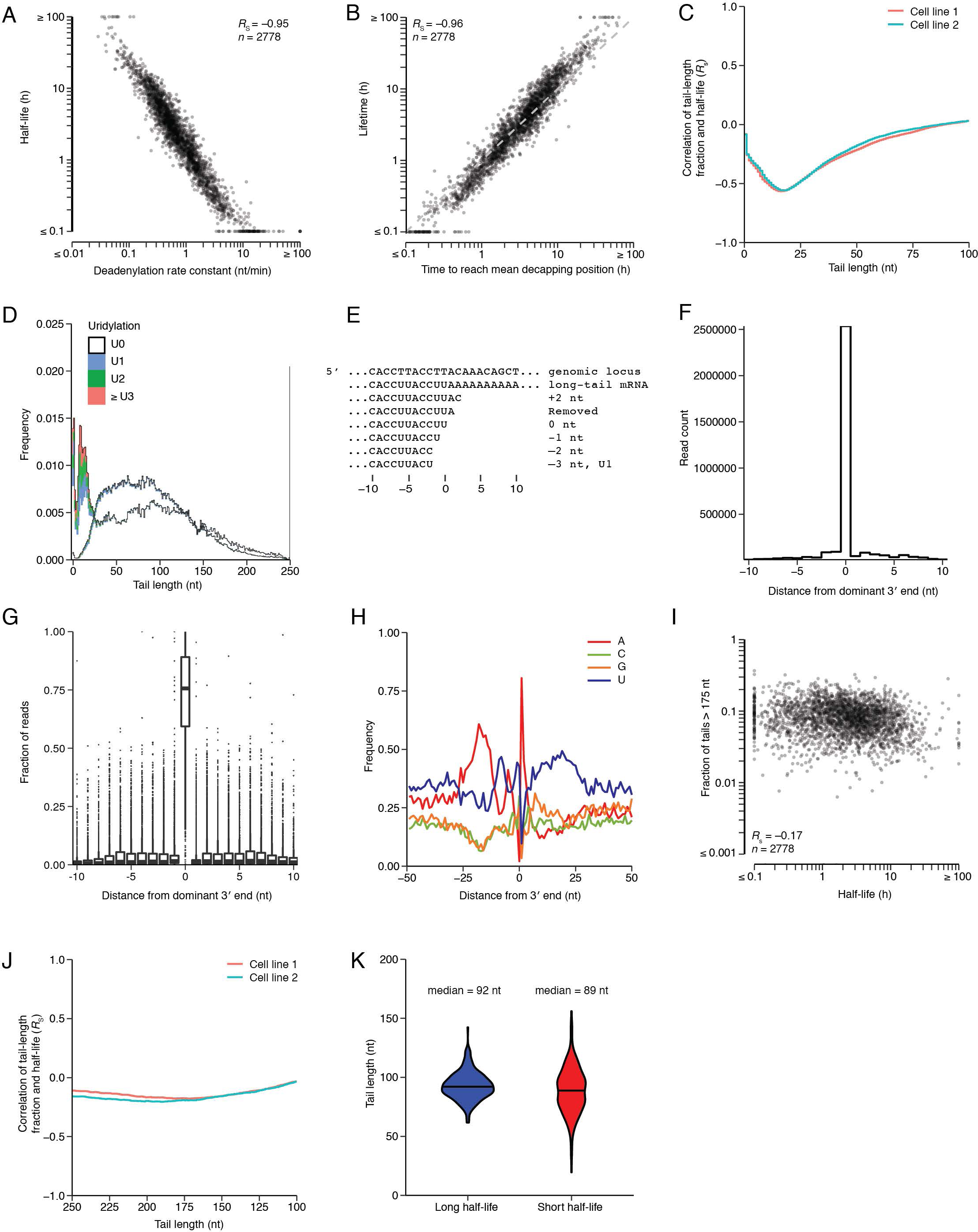
Analyses of Deadenylation Rate Constants and Steady-State Tail Lengths and Modifications, Related to Figures 5 and 6. (A) Relationship between mRNA half-life, as determined by an exponential fit to abundance, and deadenylation rate constant, as determined by the model. Because the structure of the computational model enhanced the correspondence between half-life and deadenylation rate constants, analyses performed with primary data (Figure 2B and Figure 7D) provide a more accurate indication of the correspondence between mRNA half-life and deadenlylation rate. (B) Relationship between measured mRNA lifetime and mRNA lifetime inferred by the model. For mRNAs from each gene, the number of tail nucleotides separating the mean starting tail length (Figure S4J) and the mean tail length at decapping (Figure 5C) was multiplied by the deadenylation rate constant (Figure 5A). Measured lifetime (the inverse of the degradation rate from an exponential fit) was then compared to this inferred lifetime. The dashed line indicates y = x. (C) Relationship between the steady-state abundance of short-tailed transcripts and mRNA half-life. For each tail length from 1–100 nt, the fraction of mRNAs with tail lengths that fell below that position was calculated for each gene, and the relationship (*R*_s_) between this fraction falling below the tail-length cutoff and half-life was determined as in Figure 6A. These 100 *R*_s_ values are plotted as a function of the tail-length cutoff used to classify short-tailed transcripts. This analysis started with the composite steady-state tail-length distribution generated for the analysis of Figure 2A, which accounted for very short and highly modified tails. (n = 2778). (D) The distribution of terminal uridylation on short- and long-lived mRNAs at steady state. The tail-length distributions of Figure 6B are replotted and colored by uridylation frequency (key). (E) Examples of mRNA 3′-end isoforms plotted in Figure 6E. For the genomic locus corresponding to the dominant cleavage-and-polyadenylation site of *Actb*, several possible 3′-end isoforms lacking poly(A) tails are shown, along with distance from the dominant 3′ end and whether or not the isoform would be included in the plot of Figure 6E. Also depicted is a long-tail mRNA used to annotate the 3′ end of the UTR for the analyses of Figure 6E and Figures S6F–I. (F) Distribution of tags as a function of the distance between the inferred 3′ end of their UTR and the dominant 3′ end. Only tags with a poly(A)-tail longer than 30 nt were used in this analysis. (G) Fraction of tail-containing tags for each gene as a function of the distance between the inferred 3′ end of their UTR and the dominant 3′ end. As each dominant 3′ end was defined as the position represented by the most tags in a 21 nt window, in principle, no gene should have > 50% of its tags at a position other than the dominant end. However, because dominant 3′ end annotations were determined using data from a separate experiment (standard PAL-seq with splinted ligation to the mRNA 3′ end), > 50% of tags for some genes mapped to a position other than 0; these outliers represent discrepancies between biological replicates. (H) Nucleotide composition near cleavage-and-polyadenylation sites. For each tag mapping to within 10 nt of an annotated 3′ end of an mRNA 3′ UTR, the frequency of each genomic nucleotide is plotted as a function of the distance from the annotated 3′ end. The depletion of A at position 0 and its enrichment at position 1 were artifacts of 3′-end annotation because any A at the final nucleotide of a 3′ UTR was assigned to the poly(A) tail, even if that A might have been genomically encoded. (I) Relationship between the steady-state fraction of tails > 175 nt and mRNA half-life. Otherwise as in Figure 6A. (J) Relationship between the steady-state abundance of long-tailed transcripts and mRNA half-life. For each tail length from 250–100, the fraction of mRNAs with tail lengths that fell above that position was calculated for each gene, and the relationship (*R*_s_) between this fraction falling above the tail-length cutoff and half-life was determined as in (I) and plotted as in (C). (K) Mean tail lengths for mRNAs from each gene plotted in Figure 6B. Violin plots show distributions of short- and long-lived mRNAs, with the median of each distribution shown as a horizontal line (and indicated above each group). Otherwise as in Figure 6B.

**Table S1. Parameters of the Computational Model, Related to Figure 4 and Table 1.**

Table of fitted parameters for the computational model. Staring tail lengths (*m_p_*), production rate scaling terms (*α*), deadenylation rate constants (*δ*), and decapping rate scaling terms (*β*) were fit to the computational model. The global parameters *v_p_, m_d_*, and *v_d_* were 16.27, 263.95, and 11.05 for cell line 1 and 15.5, 265.44, and 13.97 for cell line 2. mRNA half-lives were fit to an exponential model.

