## Supplemental Table 2 for "The Dynamics of Cytoplasmic mRNA Metabolism"

**Table S2. Model Parameters, Related to Figure 4**

| Parameter | Symbol | Distribution | Relevant equation | Parameter fit |
| --- | --- | --- | --- | --- |
| Production rate | *α* | Negative binomial | $k_{0}(l)= \frac{\alpha\Gamma\left( v_{p}+l \right)}{l!\Gamma\left( v_{p} \right)}\left( \frac{m_{p}}{v_{p}+m_{p}} \right)^{l}\left( \frac{v_{p}}{v_{p}+m_{p}} \right)^{v_{p}}$ | Gene specific |
| Starting tail length | *m_p_* | $''$ | $''$ | Gene specific |
| Production rate spread | *v_p_* | $''$ | $''$ | Global |
| Deadenylation | *δ* |  | $k_{1}=$*δ* | Gene specific |
| Decapping | *β* | Logistic | $k_{2}(l)= \frac{\beta}{(1+e^{-\frac{(l-m_{d})}{v_{d}}} )}$ | Gene specific |
| Mean decapping tail length | *m_d_* | $''$ | $''$ | Global |
| Decapping spread | *v_d_* | $''$ | $''$ | Global |
| The equation defining production (*k*_0_) has three fitted parameters: production rate (*α*) and starting tail length (*m_p_*), which were both fit for each gene, and spread of the starting tail length distribution (*v_p_*), which was fit globally. Deadenylation (*k*_1_) was fit for each gene as a single parameter, deadenylation rate (*δ*). Decapping (*k*_2_) uses the decapping rate (*β*) which was fit for each gene, as well as two globally fitted parameters, the decapping position (*m_d_*) and the decapping rate spread (*v_d_*). | | | | |
